# Pharmacological rescue of mitochondrial dysfunction, neurite degeneration, and premature death of ALS and AD iPSC-derived neurons

**DOI:** 10.64898/2026.04.30.722019

**Authors:** Neelam Shahani, Rupkatha Banerjee, Courtney MacMullen, Neelam Sharma, Mohammad Habibi, Henry D. Wasserman, Nathaniel C. Noyes, Puhan Zhao, Bahaa Elgendy, Michael D. Cameron, Thomas D. Bannister, Lamees Hegazy, Brian N. Finck, Ronald L. Davis

**Author notes:** Co-corresponding authors: Ronald L. Davis, Neelam Shahani.

## Abstract

Mitochondrial (MT) dysfunction is a key driver of ALS pathology. Without a healthy MT system, motor neurons (MN) function at sub-optimal levels and die. In addition, other effects of ALS, like axon/dendrite degeneration, may occur from a pathophysiological cascade spurred by MT dysfunction. A phenotypic screen identified Dipyridamole (DPM), an FDA-approved and safe drug, as having extraordinary effects on ALS patient induced pluripotent stem cell (iPSC)-derived MNs. The drug prevented MT fragmentation, loss of MT content, impaired MT bioenergetics, axon/dendrite degeneration, and premature MN death, extending neuronal survival by more than fivefold. Importantly, its efficacy extended across iPSC-derived neurons representing two different familial forms of ALS (C9orf72, TDP43) and Alzheimer’s disease (PSEN1), implying broad neuroprotection across ALS forms and other neurodegenerative diseases. DPM increased MT respiration and pyruvate uptake in a mechanism requiring the Mitochondrial Pyruvate Carrier (MPC), mechanistically explaining its biological activities. Thus, DPM is a promising drug to repurpose or refine for treating neurodegenerative diseases or other diseases that would benefit by augmenting pyruvate uptake into MT.

**Teaser:** Dipyridamole, an FDA-approved drug, restores mitochondrial function and protects neurons in ALS and Alzheimer’s disease.

## INTRODUCTION

Amyotrophic lateral sclerosis (ALS) is a fatal neurodegenerative disease characterized by progressive degeneration of motor neurons (MN) and the death of affected individuals, typically within 2–5 years after diagnosis (*1–5*). ALS represents a spectrum of diseases that includes different forms of familial ALS (fALS), familial frontotemporal dementia (fFTD), and sporadic ALS (sALS), due to different initial causes but with common neuropathology. The most common mutations leading to fALS in individuals of European descent occur in C9orf72 (34%), superoxide dismutase (SOD1, 15%), and TAR DNA binding protein (TDP43, 4%). Sporadic ALS accounts for ∼90% of cases, with susceptibility influenced by both genetic background and environmental factors. ALS neuropathology arises from complex molecular, cellular, and network-level mechanisms. Misfolded protein aggregates (SOD1, TDP43, etc.) disrupt cellular homeostasis (*6, 7*), which is exacerbated by impaired protein turnover (*8–10*). Defective nucleocytoplasmic transport, axon/dendrite degeneration, synapse loss, and glutamate excitotoxicity contribute to neuronal dysfunction (*11–15*). Oxidative stress, neuroinflammation, and altered lipid metabolism cause further neuronal damage (*6, 16–18*).

A prominent hallmark of ALS is mitochondrial (MT) dysfunction, which may explain many other pathologies through a mitochondrial cascade (*19, 20*). Abnormal MT pathology occurs early in the disease (*21–23*), supporting the idea that it triggers downstream defects. Agents that reduce MT function or content in neurites induce neurite degeneration (*24–26*). Because MT buffer intracellular Ca^2+^ (*27, 28*), their dysfunction explains excitotoxicity from glutamate-activated Ca^2+^ influx (*29*). Damaged MT are also major source of reactive oxygen species (*17, 30*), and release of MT DNA–an agonist of innate immune responses–activates inflammasomes (*31–34*). In addition, MT serve as hubs for fatty acid and lipid biosynthesis (*35*). MT localize near synapses to meet their energy demands and to provide Ca^2+^ buffering (*36*); thus MT impairment may lead to synapse loss and its consequences (*37, 38*). MT are also targets of protein aggregates that form in fALS, including ALS-C9orf72, and -TDP43 (*23, 39–43*), linking genetic insults to downstream pathologies. MT dysfunction associated with aging, acting together with genetic susceptibility or environmental factors may drive the onset of sALS. When MT become defective, opening of the MT permeability transition pore triggers apoptosis and necrosis (*44–46*). Ultimately, MT serve a key platform that determines the fate of ALS MN – cell death. The role for MT dysfunction as a main driver for ALS neuropathology predicts that drugs that protect MT from disease insults should significantly mitigate disease progression. However, no pharmacological strategy has yet been shown to robustly restore MT system performance and neuronal survival in human ALS motor neurons. We therefore asked whether pharmacologically enhancing mitochondrial function could rescue MT and neuronal phenotypes in human iPSC-derived neurons from ALS and related neurodegenerative diseases.

Here we identify the FDA-approved drug Dipyridamole (DPM) as a compound that offsets key MT, axon/dendrite, and neuronal survival phenotypes in ALS and Alzheimer’s disease (AD) iPSC-derived neurons. Mitochondrial dysfunction represents a shared pathogenic feature of both ALS and AD, suggesting that therapies targeting mitochondrial pathways may have broad relevance across neurodegenerative diseases. Phenotypic screening identified DPM as a top hit that robustly normalizes three core MT phenotypes observed in ALS and AD neurons: MT fragmentation, reduced MT number, and impaired MT bioenergetics. Our experimental approach measures MT length to report the balance between MT fission and fusion, MT count to report the balance between biogenesis and mitophagy, and MT oxygen consumption rate (OCR) to report the status of bioenergetics. DPM exhibits remarkable efficacy for normalizing MT system performance in at least two different forms of fALS, and one form of fAD. Thus, DPM exhibits efficacy in the human, disease-relevant neurons across multiple ALS forms and neurodegenerative diseases. It blocks the emergence of MT and other phenotypes in diseased iPSC-derived neurons and arrests the disease-causing progression of phenotypes after they emerge. Mechanistically, DPM enhances MT pyruvate uptake through the Mitochondrial Pyruvate Carrier (MPC), revealing a unique mechanism of action. These findings identify mitochondrial pyruvate transport as a previously underexplored therapeutic target in diseases associated with mitochondrial dysfunction.

## RESULTS

### Dipyridamole protects ALS C9orf72 lower motor neurons from mitochondrial dysfunction, axon/dendrite degeneration, and cell death

We searched for small molecules that have broad, positive effects on MT dynamics and function by screening 14,400 cpds for their effects on MT count, length, and bioenergetics, using GFP to label MT in mouse primary neurons (fig. S1) (*47, 48*). The majority of cpds selected were then screened for similar properties using doxycycline (DOX)-inducible glutamatergic neurons (i3Neurons) derived from a normal, human iPSC line (*49*), after which 17 promising cpds were subsequently screened for positive effects using diseased, iPSC-derived lower motor neurons (MN) from a patient with C9orf72 ALS (C9 MN). These funneling assays led to the identification of the well tolerated and FDA-approved drug, dipyridamole (DPM) (fig. S1).

We engineered a C9orf72 (C9) iPSC and its isogenic control line (ISO) (table S1) to stably integrate transcription factor genes (NGN2, ISL1, LHX3) at the CLYBL safe-harbor site (*50, 51*), enabling doxycycline (DOX)-inducible differentiation into lower MNs (fig. S2, A to C). After clonal selection and expansion, DOX was applied for two days to induce MN differentiation, and the resulting day 2 MN progenitors (D2 MNPs) were harvested and frozen for later replating (fig. S2, A and D). Upon replating, D2 MNPs efficiently differentiated into lower MNs, expressing key lower MN markers (HB9, ChAT, and SMI-32) as well as other neuronal markers (TUJ1, Tau, and MAP2). Notably, the ChAT-positive MNs did not express the pluripotency markers Oct4 and Sox2 (fig. S2E).

We further multiplexed our iPSC-based assays to include mScarlet as a cytoplasmic reporter (*52*), enabling segmentation and quantification of axon/dendrite (neurite) complexity (fig. S3A). Additionally, we imaged the neurons every 2 days (starting at day 8) to monitor disease progression over time, creating a multiplexed, longitudinal assay (fig. S3B). Figure S3C shows representative images from multiple days of MN differentiation using MT-GFP and cytoplasmic-mScarlet in ISO and C9-derived MN. The assay allows us to follow the onset of MT dysfunction, neurite degeneration and MN death in the same population of C9 MN as they differentiate in culture over time in comparison to ISO MN. By day 18, ALS-C9 MNs exhibit reduced MT and neurite content compared to ISO MNs, and dead neurons; small, circular MT remnants; and neurite debris (fig. S3C, day 18 and inset, images within the red box). Our multiplexed, longitudinal, and phenotypic assay using iPSC-derived neurons thus provides metrics for multiple phenotypes that occur in ALS MN, including MT count (average number of MTs), MT content (cumulative area, or CA, representing the pixel area occupied by MTs in each image), MT length (average MT length), neurite CA (cumulative area occupied by neurites, serving as a measure of neurite integrity/degeneration), and MN survival (fig. S3C). These metrics together provide measures for ALS MN pathology, including MT fragmentation, loss of MT content, neurite degeneration, and MN survival.

Fig. 1, A and B present the assay timeline and quantitative data for these metrics using C9 and ISO iPSC-derived MNs. For DMSO-treated (on day 12) control ISO MN, we observed a progressive increase in MT count, CA, length, and neurite CA over time in culture, providing the baseline for MT health, neurite integrity and survival of control ISO MNs (Fig. 1B). For DMSO-treated C9 MN, the MT and neurite parameters were similar to ISO MNs from days 8 to 12 but began a significant and progressive decrease starting on day 14/16, indicating progressive deterioration of MT and cell health. Thus, the longitudinal assay of human, ALS-diseased iPSC-derived MN captures the progressive MT, axon/dendrite, and poor survival pathology observed in ALS postmortem tissue and animal models.

**Fig. 1:**
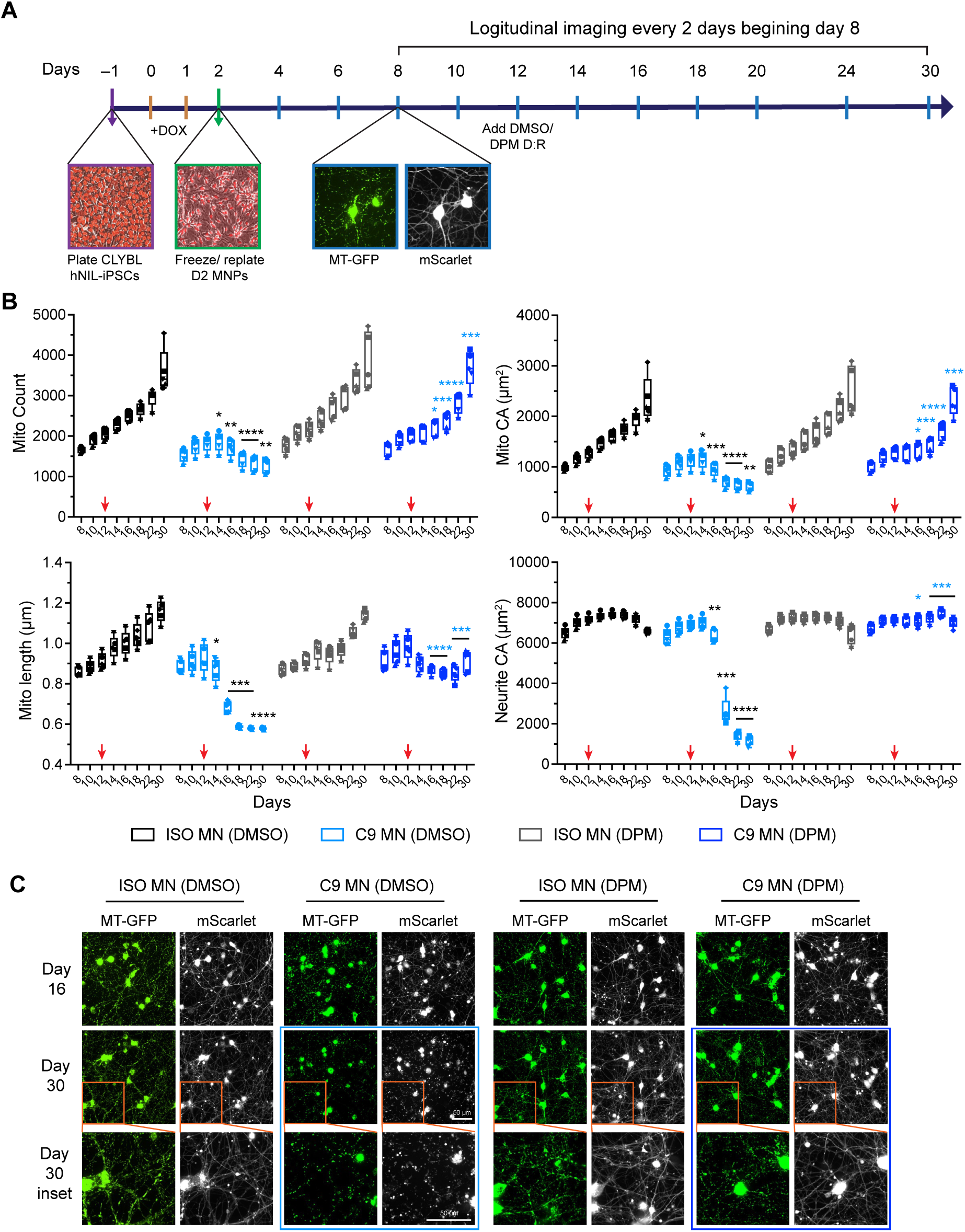
DPM is neuroprotective for C9 iPSC-derived MNs. **(A)** Overview of the MT/neurite dynamics assay time-course from plating CLYBL hNIL iPSCs through MN neuron differentiation, MN culture with mtTagGFP2-mScarlet lentivirus at the appropriate MOI, and longitudinal imaging every two days beginning day 8. DPM/DMSO addition was performed at day 12. **(B)** Strong, reproducible, and progressive MT (count, CA, length) and neurite CA phenotypes of C9 MN compared to the ISO MN across days in culture in the time-course assay treated with single dose of DMSO or DPM. CA=cumulative area, the number of pixels representing MT or neurites in each image. Data are presented as box plots with individual data points (unique symbols) from n = 5 independent time-course experiments. The box extends from the 25th to 75th percentiles, with the line in the box indicating the median. The whiskers extend to the minimum and maximum values. Within each independent experiment, the DMSO group included n = 40 technical replicates, while the DPM group included n = 5 technical replicates. Red arrows indicate the addition of single dose of DMSO (0.125%) / DPM (10 µM) on day 12. Black asterisks illustrate the significant differences of the C9 MN (DMSO) vs ISO MN (DMSO). Blue asterisks illustrate the significant differences of the C9 MN (DPM) vs C9 MN (DMSO). *p<0.05, **p<0.01, ***p<0.001, ****p<0.0001, Two-way repeated measures ANOVA with Tukey’s multiple comparisons test. **(C)** Representative images of C9 MN treated with DMSO or DPM at day 12 and imaged at day 16, and day 30. Day 30 inset (orange box) indicate magnified images. Healthy DPM-treated neurons (dark blue box around images, d30 and d30 inset) compared to the degenerated DMSO-treated C9 MN (light blue box around the images showing small fragments of MT and cell debris).

The C9 and ISO MNs were treated with a single dose of DPM, in dose response (D:R), at day 12 before the MT pathology becomes apparent. This early treatment significantly protected MT and neurite health in the C9 MNs, evident by the normalization of MT count, MT CA, and neurite CA, and partial normalization of MT length in C9 MNs compared to the ISO MNs (Fig. 1B). Surprisingly, DPM had no significant effect on the ISO MN. Similar results were obtained from collapsing the data from five independent experiments (Fig. 1B) or from a single experiment (fig. S4A), demonstrating the assay’s reproducibility in detecting MT and neurite phenotypes and the neuroprotective effects of DPM. Dose response data collected for the metrics on each imaging day show that DPM exhibits potency (EC_50_) in the low micromolar range and a very high efficacy (span) (fig. S4B and table S2).

Images shown in Fig. 1C illustrate that MT and neurites in C9 MNs maintained a healthy appearance at day 30 when treated with DPM; whereas C9 MNs treated with DMSO displayed degeneration with many small MT fragments and cell debris (Fig. 1C, day 30 inset). This degeneration correlates with the floor values for the MT/neurite metrics reached around day 18 (Fig. 1B), indicating that the metrics at ∼day 18 mark the time of death of most or all MNs, which correlates with a visual determination. We quantified neuroprotection using the RealTime-Glo Cell Viability reagent at day 31 to both ISO and C9 MNs with DPM in D:R and analyzing the results on day 33. The EC_50_ for this assay was in the low micromolar range (fig. S5A). Remarkably, a single treatment with 10 µM DPM significantly extended the survival of C9 MN from day 18 to more than day 100 (fig. S5B and movie S1). Some of DPM’s remarkable and persistent effects can be explained by its metabolic stability in culture. Measurements of DPM in the media of cultured MN indicate that it is slowly metabolized with a half-life of approximately 9 days (fig. S5, C and D).

We wondered if DPM would offer neuroprotection when provided after the onset of neuropathology in the C9 MN. To test this, we provided a single dose of DPM at various days to cultures of C9 MN. For cultures receiving DPM on day 12, MT count, MT CA, and neurite CA differed from the DMSO groups starting at day 18, while MT length differed at day 16 (Fig. 2). But irrespective of whether DPM was administered at day 16, 18, or 20, the drug provided significant protection at subsequent days for all four metrics, indicating that DPM both arrests the progression of pathology and protects from its onset. We also found that earlier administration of DPM at day 8 produced a significant increase in MT CA and MT length along with improved EC_50_ and span values compared to the day 12 addition (fig. S4, C and D; table S2). Thus, earlier intervention with DPM enhances its neuroprotective effects, leading to more favorable MT dynamics and improved neuronal health.

**Fig. 2:**
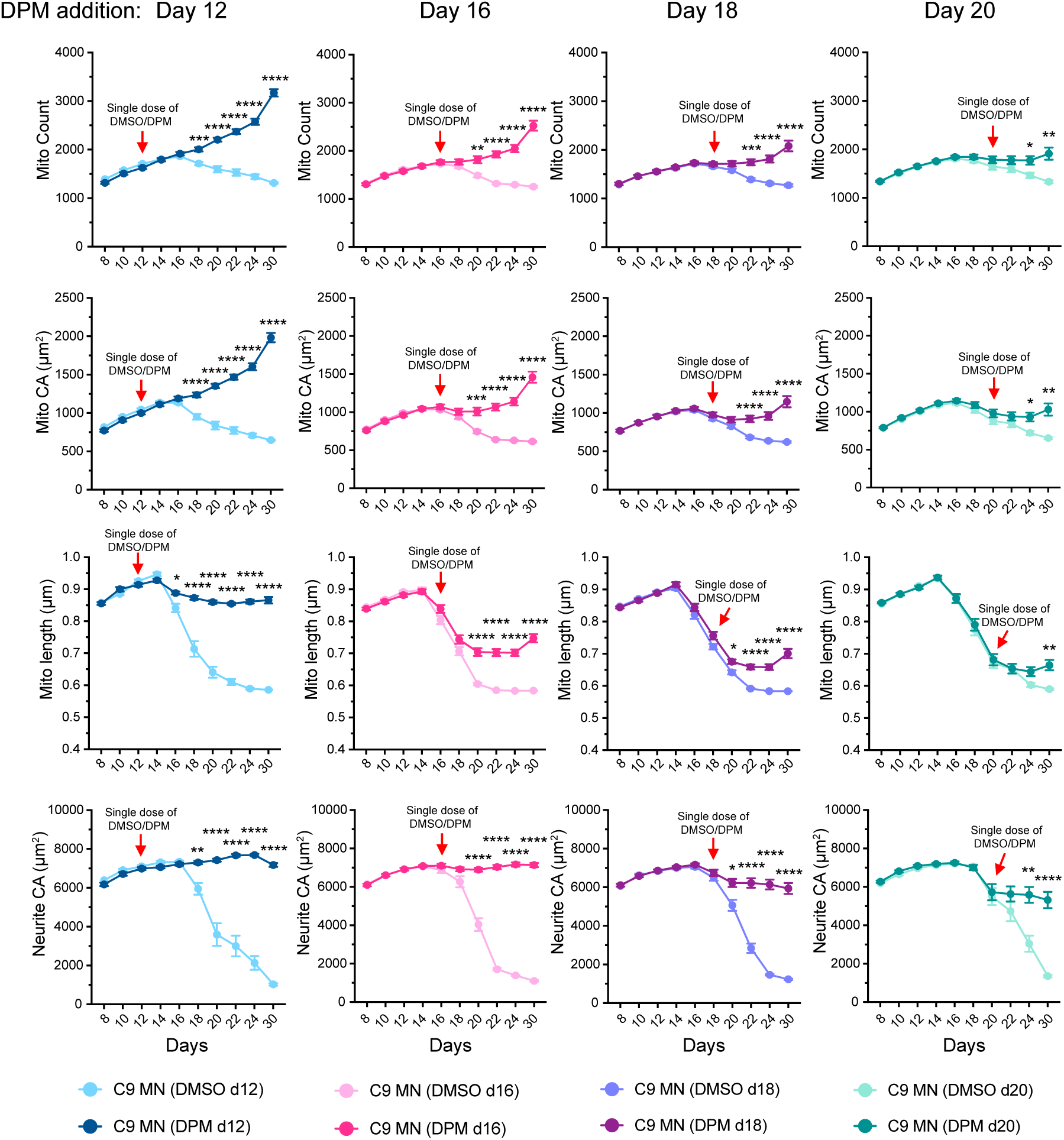
DPM arrests the progression of MT and neurite phenotypes in C9 MN. Time-course experiment as in Fig. 1, with single dose of DPM added before pathology onset on day 12, or after onset on days 16, 18, or 20 to C9 MN. Red arrows indicate the addition of single dose of DMSO (0.125%) / DPM (5 µM) after the imaging on the respective days. Data are mean ± SEM. *p<0.05, **p<0.01, ***p<0.001, ****p<0.0001, Two-way ANOVA, with Šídák’s multiple comparisons test (n = 22-24 technical replicates for DMSO and n = 22-24 technical replicates for DPM-treated groups).

The prior experiments were performed using C9 and ISO MN derived from iPSCs sourced from Cedar Sinai. To ensure that the results obtained were not specific to this one C9/ISO iPSC pair, we tested the effects of DPM using three other C9 iPSC lines from different ALS patients (table S1). Although the time-course trajectory of the metrics measured differ across the lines – probably due to genetic background and most notably for 33.1 C9 MN – the results show that a single dose of DPM added on day 12 significantly increased MT and neurite parameters across all lines (fig. S6). This indicates that the neuroprotective effects of DPM are not limited to a single patient-specific model but have broader relevance across various genetic backgrounds.

We next used the Seahorse XF Mito Stress Test to assess key parameters of MT function by directly measuring the oxygen consumption rate (OCR) of MN. Pilot experiments revealed deficits in OCR parameters as early as day 14, so we performed a subsequent D:R experiment, with DPM added on day 8. DPM significantly protected the MT function in the diseased C9 MN, improving all four standard respiratory parameters in a dose-dependent manner to levels like those of ISO MN (Fig. 3, A and B). Notably, DPM’s EC_50_ values for these MT functional parameters were in the sub-micromolar range (Fig. 3C), indicating its high potency in restoring MT function. These findings align with the improvements observed in MT dynamics when DPM was added earlier on day 8 and show that DPM’s protective effects extend to MT bioenergetics in C9 MNs.

**Fig. 3:**
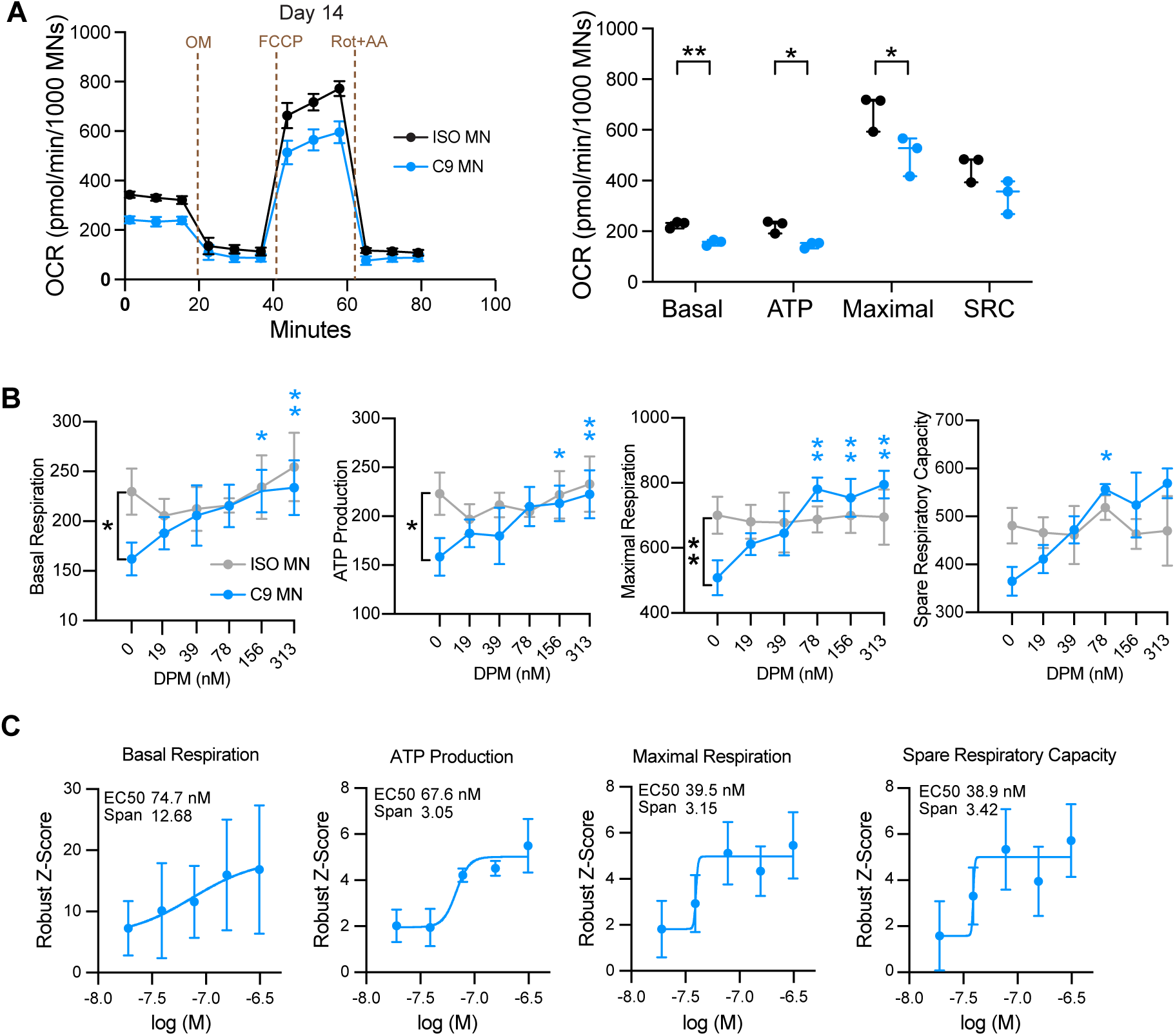
DPM rescues MT bioenergetics deficits in C9 MN. **(A)** MT function was assessed in ISO and C9 MN at day 14 by measuring OCR using the Seahorse XFe96 Analyzer. The Mito Stress Test was used to measure bioenergetics parameters, by adding sequentially the Complex V inhibitor oligomycin (OM), mitochondrial uncoupler carbonyl cyanide-p-trifluoromethoxyphenylhydrazone (FCCP), and Complexes I and III inhibitors rotenone (Rot) and antimycin A (AA). Basal respiration, ATP-linked respiration, maximal respiration, and spare respiratory capacity (SRC) were calculated, and OCR values were normalized to live MN counts/well (from 4 fields/well) determined using Calcein AM staining and presented as pmol/min/1000 MNs. Data were analyzed by comparing the average of 15 technical replicates across 3 independent experiments. *p<0.05, **p<0.01 unpaired Student’s t-test. **(B)** DPM at indicated concentrations was added on day 8 and the Mito Stress Test was performed on day 14. Black asterisks illustrate the significant difference in C9 MN vs ISO without DPM, and blue asterisks illustrate the significant difference in C9 MN with DPM vs C9 MN with no DPM. Data are mean ± SEM. *p<0.05, **p<0.01, Two-way ANOVA, Šídák’s multiple comparisons test (n = 3 independent experiments with 8 technical replicates per experiment). **(C)** D:R curves from (b) to obtain EC_50_ and span values for each MT respiratory parameter: basal respiration, ATP-linked respiration, maximal respiration, and spare respiratory capacity. D:R data (mean of 3 independent experiments) are presented as robust Z-scores for DPM-treated C9 MN relative to in-plate DMSO control wells. Robust Z-scores ≥2 or ≤- 2 are considered significant. Potency is measured by EC_50_ and efficacy by span.

Together, these data show that DPM robustly protects C9 MN from five different ALS phenotypes: MT fragmentation, loss of MT content, and impaired bioenergetics; along with axon/dendrite degeneration and early MN death. In contrast, riluzole and edaravone, two FDA-approved drugs for ALS treatment, exhibited little efficacy in our longitudinal assay and neither drug extended MN survival (fig. S7).

### Dipyridamole provides neuroprotection to TDP43 ALS MN and Alzheimer’s disease cortical neurons

We then wondered whether DPM might provide similar protection to neurons representing other forms of fALS and other forms of neurodegenerative disease. We first tested DPM in MN generated from iPSCs heterozygous (HET) for an engineered, ALS-causing TDP43^M337V^ mutation in a normal control genetic background. As the control, we used MN made from iPSCs engineered to revert (REV) the mutation back to the wildtype allele (Kolf2.1J, (*53*)). We observed a significant and progressive decrease in the MT count, MT CA, MT length and neurite CA over time in culture for the TDP43^M337V^ MNs, starting at day 14, compared to the control REV MNs (Fig. 4A). MT length for the TDP43^M337V^ MNs reached floor values by day 20, indicating severe deterioration of health. REV MN were healthier but also showed a decrease in the MT and neurite parameters at the later time points. The decreases using the REV MN are in contrast with the ISO C9 MN, suggesting a significant influence of genetic background on neuronal health. Nevertheless, a single dose of DPM at day 10 increased the MT and neurite metrics in both TDP43^M337V^ MN and REV MN (Fig. 4A), with EC_50_s for TDP43^M337V^ MN in the low micromolar range (Fig. 4B and table S2). These results indicate that DPM treatment protects against MT and neurite degeneration and enhances neuronal survival in both diseased and control neurons. Additionally, DPM improved MT bioenergetics deficits in TDP43^M337V^ MN (Fig. 4, C and D), although with 2-3X higher EC_50_s compared to C9 MN, suggesting that while effective, DPM may require higher concentrations to achieve similar potency in the TDP43 MN. Most importantly, the results show that DPM is effective in at least two distinct fALS MN models, suggesting that it may be effective across multiple forms of ALS.

**Fig. 4:**
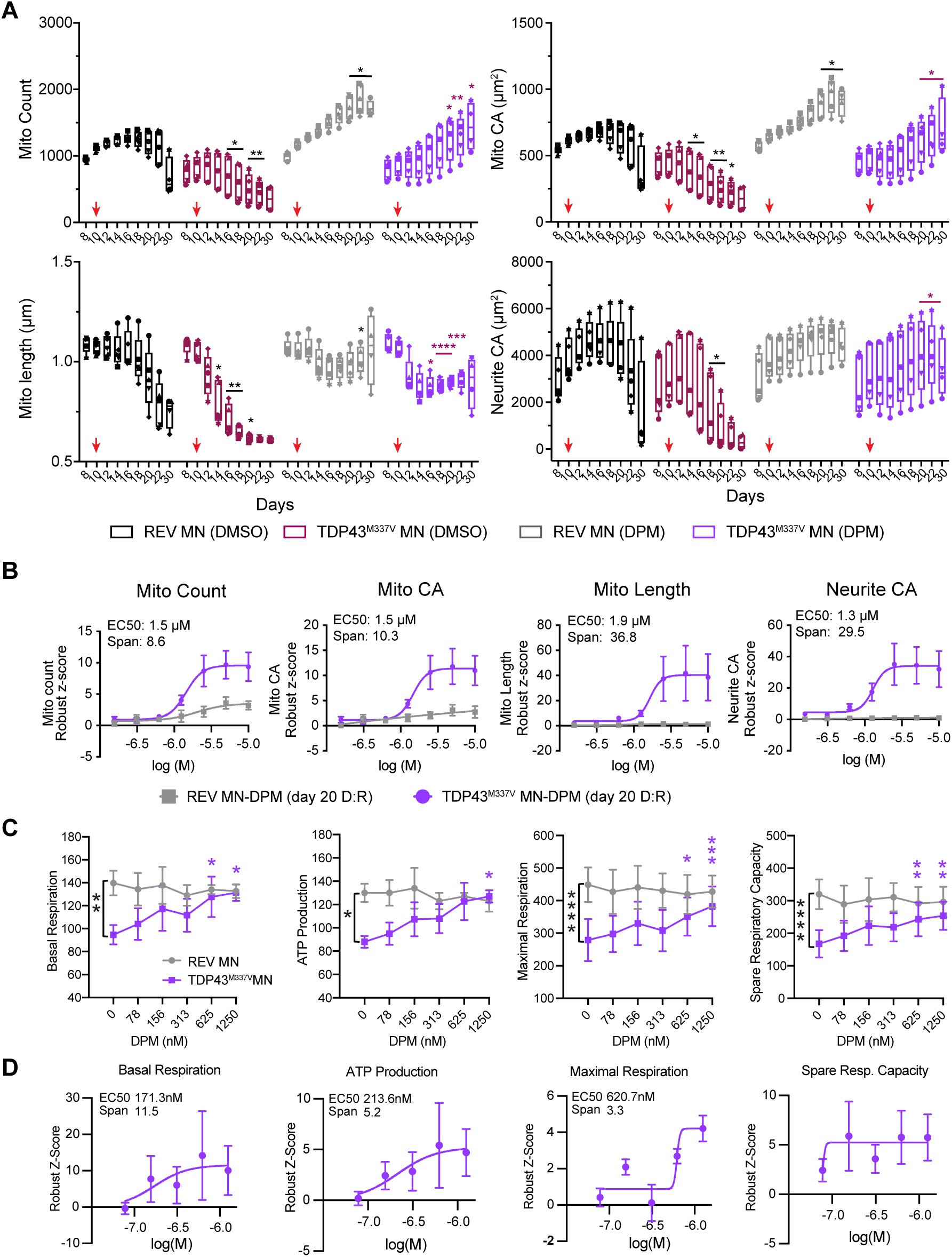
DPM is neuroprotective and improves MT bioenergetics deficits in TDP43^M337V^ iPSC-derived MNs. **(A)** Progressive MT (count, CA, length) and neurite (CA) phenotypes in TDP43^M337V^ MN compared to the REV MN across days in culture in the time-course assay treated with single dose of DMSO or DPM. Data are presented as box plots with individual data points (unique symbols) from n = 4-5 independent time-course experiments. Box plots show the median (center line), the 25th to 75th percentiles (box), and minimum and maximum values (whiskers). Within each experiment, the DMSO group includes n = 6-9 technical replicates, while the DPM group includes n = 5-11 technical replicates. Red arrows indicate the addition of single dose of DMSO (0.125%) / DPM (10 µM) on day 10. Black asterisks illustrate the significant differences of the TDP43^M337V^ MN (DMSO) vs REV MN (DMSO) and REV MN (DMSO) vs REV MN (DPM). Maroon asterisks illustrate the significant differences of the TDP43^M337V^ MN (DPM) vs TDP43^M337V^ MN (DMSO). *p<0.05, **p<0.01, ***p<0.001, ****p<0.0001, Two-way repeated measures ANOVA with Tukey’s multiple comparisons test. **(B)** D:R curves for DPM at day 20 using TDP43^M337V^ MN and REV MN. D:R data (mean ± SEM of 5 independent experiments) for MT count, MT CA, MT length, and neurite CA are presented as robust Z-scores using TDP43^M337V^ MN / REV MN relative to the respective in- plate DMSO control wells. DPM was added on day 10 and tested from 156 nM-10,000 nM. The measured EC_50_ values and spans for the four parameters using the TDP43^M337V^ MN are shown. **(C)** DPM at indicated concentrations was added on day 8 and the Mito Stress Test was performed on day 14 as in Fig. 3. Black asterisks illustrate the significant difference in TDP43^M337V^ MN vs REV without DPM, and purple asterisks illustrate the significant difference in TDP43^M337V^ MN with DPM vs TDP43^M337V^ MN with no DPM. Data are mean ± SEM. *p<0.05, **p<0.01, ***p<0.001, Two-way ANOVA, Šídák’s multiple comparisons test (n = 3 independent experiments with 8 technical replicate per experiment). **(D)** Dose response (D:R) curves from (c) to obtain EC_50_ and span values for each MT respiratory parameter: basal respiration, ATP-linked respiration, maximal respiration, and spare respiratory capacity. D:R data (mean of 3 independent experiments) are presented as robust Z-scores for DPM-treated C9 MN relative to in-plate DMSO control wells.

Given the neuroprotection offered to iPSC-derived MN representing different forms of ALS, we hypothesized that DPM might exhibit efficacy to iPSC-derived neurons representing other types of neurodegenerative diseases. To test this possibility, we differentiated iPSC-derived glutamatergic cortical neurons (CN) carrying the Alzheimer’s disease PSEN1 A246E mutation (AG67) together with their isogenic control (ISO-AG67) (*52*). We observed a highly significant and progressive decline in MT length and a more modest decline in MT count in AG67 neurons compared to ISO-AG67 (Fig. 5A). By day 22, both ISO-AG67 and AG67 neurons exhibited deteriorating health, with complete neuronal death occurring by day 26, as indicated by MT/neurite metrics reaching floor values. We observed that a single dose of DPM administered on day 10 significantly improved MT health for both ISO-AG67 and AG67 neurons but had a limited impact on survival (Fig. 5A), with D:R curves (Fig. 5B and table S2) revealing more limited efficacy compared to the ALS C9 MN. Nevertheless, DPM improved MT bioenergetics deficits in AG67 CN (Fig. 5, C and D), with EC_50_s in low micromolar range. The overall data indicate that DPM significantly offsets the MT pathology and bioenergetics deficits in the PSEN1 CN, although the efficacy is reduced relative to its effects in ALS MN.

**Fig. 5:**
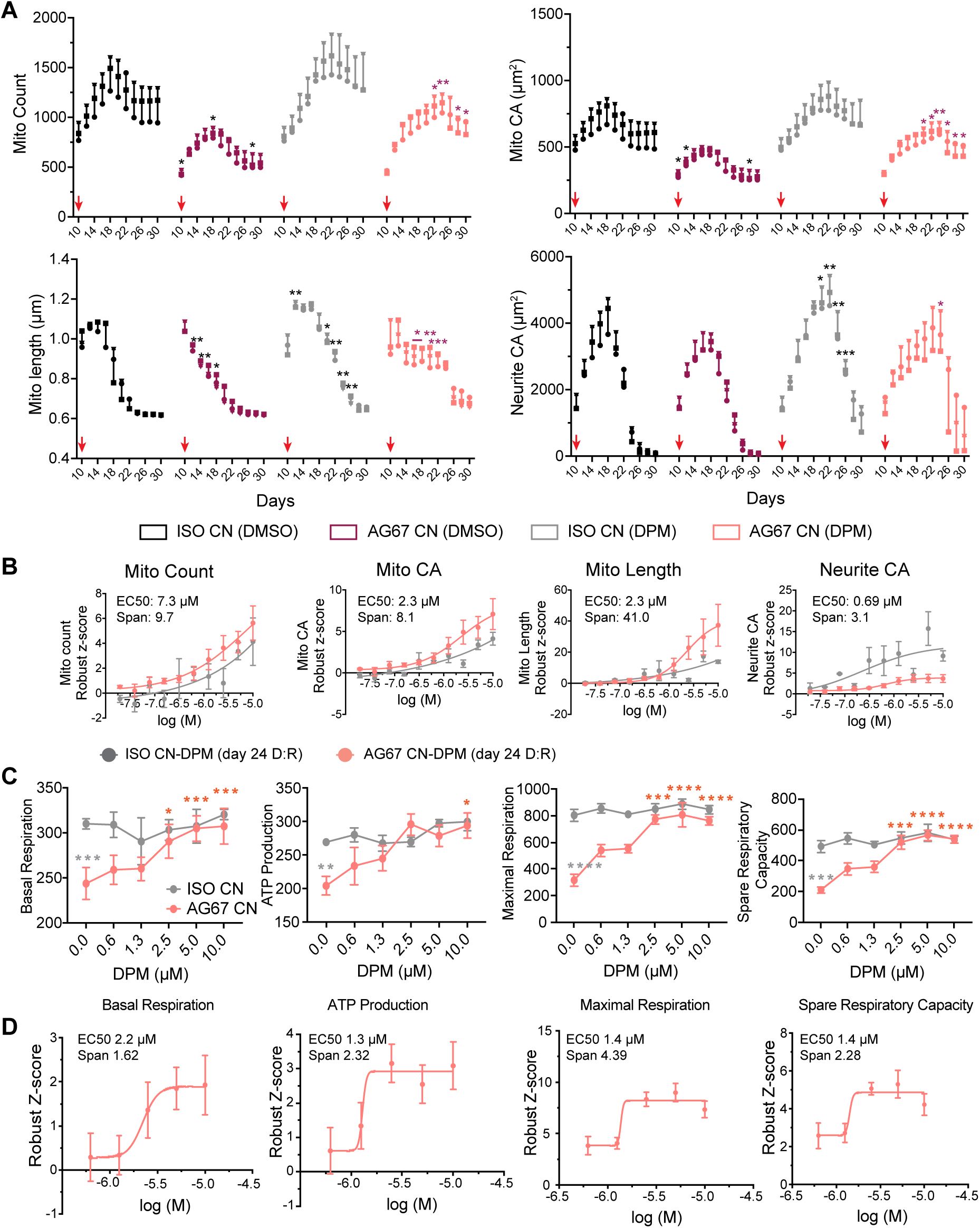
DPM is neuroprotective and improves MT bioenergetics deficits in PSEN1^A246E^ cortical neurons. **(A)** MT (count, CA, length) and neurite CA phenotypes in AG67 (PSEN1^A246E^) cortical neurons (CN) compared to the ISO AG67 CN across days in culture in the time-course assay. AG67 and ISO CN were imaged every other day beginning at day 10. A single dose of DPM and/or DMSO was added on day 10 after imaging. Data from three independent experiments are depicted with each unique symbol point representing the average of each parameter at each time point. Black asterisks illustrate significant differences comparing AG67 CN (DMSO) vs ISO CN (DMSO) and ISO CN (DPM) vs ISO CN (DMSO). Maroon asterisks illustrate significant differences between AG67 CN (DPM) vs AG67 CN (DMSO). *p<0.05, **p<0.01, ***p<0.001, Two-way repeated measures ANOVA with Tukey’s multiple comparisons test (n = 3 independent time course experiments) with n = 40 replicates (DMSO group) and n = 5 replicates (DPM group) in each experiment. Red arrows indicate the addition of single dose of DMSO (0.125%) / DPM (5 µM) on day 10. (**B**) D:R curves for DPM at day 24 using AG67 CN and ISO CN. D:R data (mean ± SEM of 3 independent experiments) for MT count, MT CA, MT length, and neurite CA are presented as robust Z-scores using AG67 CN / ISO CN relative to the respective in-plate DMSO control wells. DPM was added on day 10 and tested at concentrations ranging from 156 nM-10,000 nM. The measured EC_50_ values and spans for the four parameters are shown. **(C)** DPM at indicated concentrations was added on day 8 and the Mito Stress Test was performed on day 16 as in Fig. 4. The DPM D:R was normalized to Calcein AM live cell fluorescence intensity (from whole well). Grey asterisks illustrate the significant difference in AG67 CN vs ISO CN without DPM, and red asterisks illustrate the significant difference in AG67 CN with DPM vs AG67 CN with no DPM. Data are mean ± SEM. *p<0.05, **p<0.01, ***p<0.001, ****p<0.0001, Two-way ANOVA, Šídák’s multiple comparisons test (n = 3 independent experiments with 6 technical replicate per experiment). **(D)** D:R curves from (c) to obtain EC_50_ and span values for each MT respiratory parameter: basal respiration, ATP-linked respiration, maximal respiration, and spare respiratory capacity. D:R data (mean of 3 independent experiments) are presented as robust Z-scores for DPM-treated AG67 CN relative to in-plate DMSO control wells.

### Dipyridamole does not exert its effects by inhibiting PDEs, ENTs or SREBPs

DPM is used clinically as an anticoagulant, commonly administered after heart surgery to reduce the risk of blood clots and as a preventive treatment for stroke (*54*). Two mechanisms of action account for this activity: inhibition of cyclic nucleotide phosphodiesterases (PDEs) and equilibrative nucleoside transporters (ENT) 1 and ENT2 (*55–58*). We tested whether these actions are responsible for its remarkable ability to enhance MT health and viability in ALS and AD neurons, by assaying the effects of 12 PDE and 3 ENT inhibitors in dose:response in our longitudinal assay to see if they mimic DPM’s effects on MT health, neurite complexity, and MN survival. We observed that none of the inhibitors tested showed positive effects in the assay (fig. S8, A, B, and D; tables S3 and S4), except for the non-selective PDE inhibitor Zaprinast (see below). DPM also suppresses SREBP activation independently of its PDE inhibition, suggesting a possible role in lipid regulatory pathways (*59, 60*). However, 3 selective SREBP inhibitors failed to replicate DPM’s protective effects (fig. S8, C and D; table S5). Together, these results indicate that DPM’s neuroprotective effects on MT health and neuronal viability are mediated by a mechanism distinct from its known inhibition of PDE, ENT, and SREBP pathways.

### Dipyridamole increases pyruvate access into mitochondria through the mitochondrial pyruvate carrier

Of the 12 PDE inhibitors we tested, only a single dose of Zaprinast added on day 12, significantly protected against MT and neurite degeneration and improved neuronal survival in C9 MN. However, this protection was limited to day 24, with EC_50_s in low micromolar range comparable to DPM (fig. S9 and table S3). Notably, Zaprinast has also been shown to inhibit the mitochondrial pyruvate carrier (MPC) (*61, 62*), prompting us to explore whether DPM might act through a related mechanism involving pyruvate access to MT through MPC function.

We first performed in silico docking studies of DPM to the crystal structure of the MPC. The MPC is a functional heterodimer consisting of the MPC1 and MPC2 proteins each containing three transmembrane domains and N-terminal amphipathic helices containing hydrophobic residues allowing them to lie on the inner mitochondrial membrane (*63*). However, there remains controversy from four recently published cryoEM studies regarding MPC orientation, with the structurally more open end proposed to be oriented towards the matrix (matrix open) in one study and towards the intermembrane space (IMS open) in the three other studies (*64–67*).

Fig. 6A illustrates docking poses of DPM within both matrix-open and IMS-open conformations of the MPC heterodimer. In the matrix-open conformation (Fig. 6A, left) (PDB: 9MNW), DPM exhibited a predicted binding energy of –6.6 kcal/mol, which was weaker than that of the reference inhibitor GW604714 (–8.7 kcal/mol). In this binding mode, key residues for binding included P30, N77, V73, H26, F66 and F69 from MPC1, and W82, N100, Y85, F42, I89, N93, S86, and L96 from MPC2. The relatively large size of DPM may hinder its translocation through the MPC channel and alternatively diffuse across the inner mitochondrial membrane (IMM) into the matrix to access the binding pocket, as proposed for inhibitors such as UK-5099, AKOS, and GW604714 (*64*). Fig. 6A (right) illustrates the IMS-open conformation (PDB: 9MNY). DPM does not fit optimally into the canonical pyruvate-binding pocket. However, docking identified a secondary, druggable site toward the IMS, where DPM achieved a binding energy of –5.6 kcal/mol. In this site, the ligand formed a balanced network of interactions, engaging in hydrogen bonding with polar residues at the cavity entrance while positioning its hydrophobic regions against nonpolar side chains deeper within the pocket. Binding was mediated by MPC1 residues M55, L36, N33, and Y62, and MPC2 residues W74, R62, K49, T78, V103, A106, L75, Q71, M60, D59, and V53. This pose provided better steric complementarity than the pyruvate pocket in the matrix-open state, though the overall binding energy was still lower than the top-scoring matrix-open pose. This alternative site may afford the identification of MPC modulators with a novel mechanism of action.

**Fig. 6:**
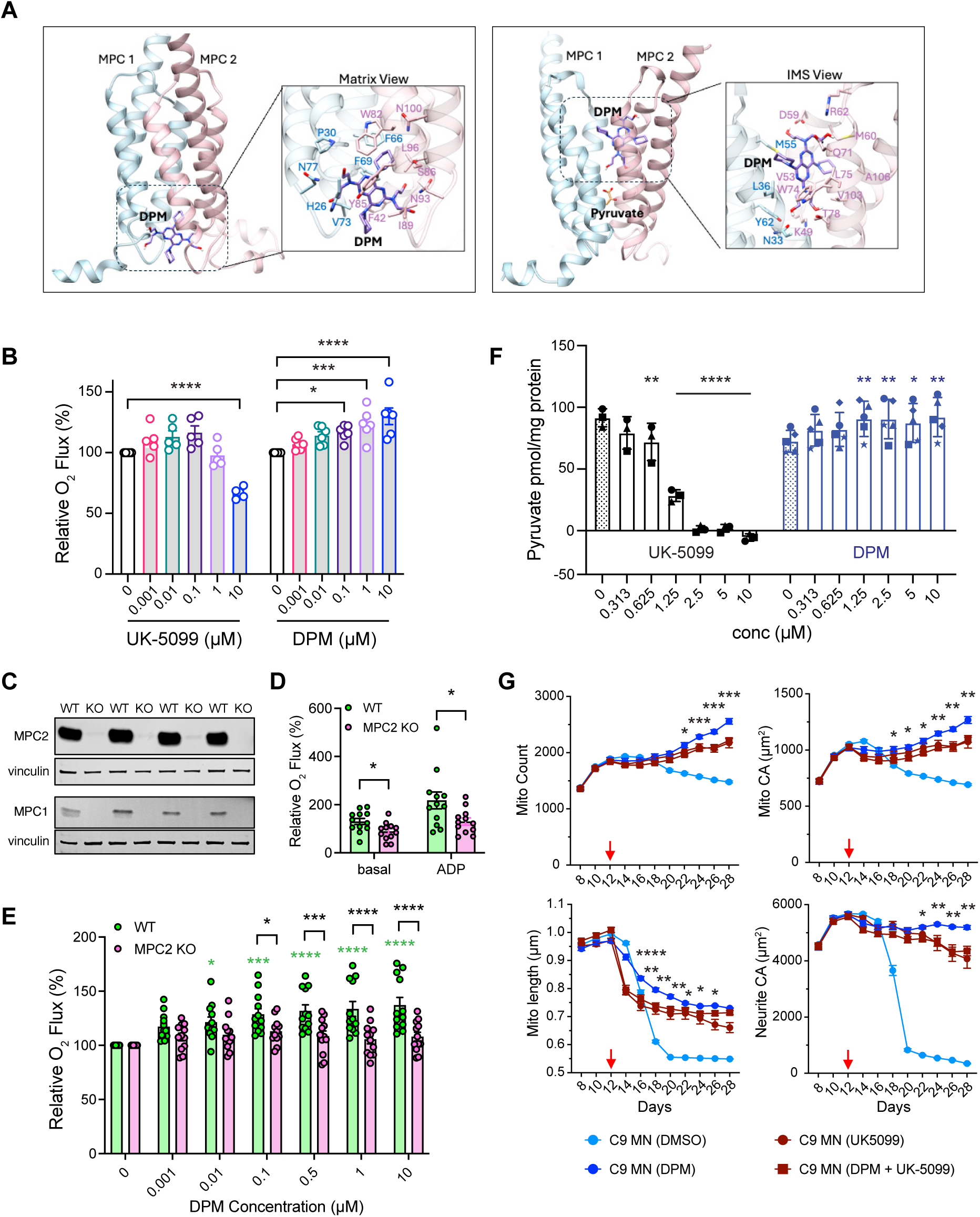
DPM increases pyruvate access to the mitochondria through the mitochondrial pyruvate carrier. **(A)** Left: Docking pose of DPM in the matrix-open conformation of the MPC heterodimer (PDB: 9MNW). MPC1 is depicted as blue ribbons and MPC2 as red ribbons. Interacting residues are displayed in stick representation, and DPM is shown as purple sticks. Right: Docking pose of DPM in the IMS-open conformation of the MPC heterodimer (PDB: 9MNY). **(B)** Oxygen (O_2_) flux of isolated mouse liver MT in response to increasing concentrations of UK-5099 or DPM. For each technical replicate, baseline respiration (0 µM; ADP, pyruvate, and malate present) was set to 100%, and O_2_ flux following compound addition was normalized to this baseline. Data are mean ± SEM. *p<0.05, ***p<0.001, and ****p<0.0001 versus baseline values. Two-way ANOVA with Tukey’s multiple comparisons test [n = 5 (UK-5099) or n = 6 (DPM) samples from 3 independent experiments]. **(C)** MPC1 and MPC2 protein expression assessed by Western blots in liver MT from four mice of each genotype [wildtype (WT) and MPC knockout (KO)] used for O_2_ flux assays shown in panels D and E. **(D)** Basal and ADP-stimulated O_2_ flux of isolated liver MT from WT and MPC2 KO mice shown in panel C, plotted without normalization. This illustrates the significantly lower baseline respiration observed in MPC2 KO MT compared with WT. Data are mean ± SEM. *p<0.05, two-sided Student’s *t*-test. **(E)** O_2_ flux of isolated liver MT from WT and MPC2 KO mice in response to increasing concentrations of DPM. For each technical replicate, baseline (0 µM DPM; ADP, pyruvate, and malate present) was set to 100%, and O_2_ flux following DPM addition was normalized to this baseline. Data are mean ± SEM. Green asterisks indicate significant differences between DPM concentrations and the baseline (0 µM) in WT MT. Black asterisks indicate significant differences between WT and MPC2 KO MT at the indicated DPM concentrations. *p<0.05, ***p<0.001, and ****p<0.0001 versus baseline values. Two-way ANOVA with Tukey’s multiple comparisons test [n = 3 (DPM) samples from 4 independent experiments for each genotype]. **(F)** MT pyruvate uptake assay using mouse liver MT using increasing concentrations of UK-5099 (n=3 independent experiments) and DPM (n=5 independent experiments). Data are mean ± SEM. *p<0.05, **p<0.01, ****p<0.0001, mixed-model ANOVA followed by student’s *t*-test using the standard least squares model in JMP v15. **(G)** MPC inhibition is dominant over the augmenting effects of DPM on MT and neurite integrity in C9 MN. Longitudinal assays as in Fig. 2 using single doses of 10 µM DPM and 5 µM UK-5099 added before pathology onset on day 12. Co-treatment with UK-5099 overrides the effects of DPM, consistent with the MPC-dependent mechanism observed with isolated MT shown in panel E. Red arrows indicate the addition of single dose of DMSO (0.125%) / DPM (10 µM) / UK-5099 (5 µM) / DPM (10 µM) + UK-5099 (5 µM) on day 12. Asterisks indicate significant differences between DPM alone and the DPM+UK-5099 conditions. Data are mean ± SEM. *p<0.05, **p<0.01, ***p<0.001, ****p<0.0001, Two-way repeated measures ANOVA with Tukey’s multiple comparisons test (n = 36 technical replicates for DMSO and n = 11 technical replicates for DPM, UK-5099, and DPM+UK-5099 groups).

The suggested physical interaction between DPM and MPC led us to test whether DPM directly influences MT pyruvate metabolism. MT were isolated from wildtype mouse liver and rates of oxygen flux in the presence or absence of DPM were assessed by using high resolution respirometry. As expected, incubating mitochondria with UK-5099, a known MPC inhibitor, suppressed pyruvate dependent respiration at 10 µM. But to our surprise, incubation with 0.1, 1, or 10 µM DPM significantly enhanced pyruvate-mediated respiration by up to ∼30% in a dose dependent manner (Fig. 6B). To determine whether this effect required MPC, we repeated the experiment using MT isolated from the livers of MPC2 knockout mice, which lack both MPC1 and MPC2 proteins (Fig. 6C) since only the heterodimer is stable. The MPC1/MPC2 knockout MT displayed reduced baseline respiration (Fig. 6D), and consistent with our initial observations, DPM enhanced respiration in wildtype MT in a dose dependent manner. Importantly, this effect was completely abolished in MPC-deficient MT (Fig. 6E). These data suggest that DPM augments – in an MPC-dependent manner – rather than inhibits, pyruvate metabolism in isolated MT, as reflected by increased oxygen flux.

We tested the hypothesis that DPM enhances pyruvate uptake directly using a modified radiometric assay for MT pyruvate uptake and mouse liver MT (*68*). As expected, UK-5099 inhibited ^14^C-pyruvate uptake in a dose dependent manner (Fig. 6F). In contrast, DPM significantly augmented ^14^C-pyruvate uptake by up to ∼22% at concentrations ranging from 1.25-10 µM, consistent with the results obtained from respirometry, and leading to the conclusion that DPM enhances pyruvate uptake by MT through MPC function.

Given the surprising but opposing effects of UK-5099 and DPM on pyruvate uptake using isolated MT, we performed a competition experiment to evaluate their functional interaction in our longitudinal MT dynamics assay using C9 MN (Fig. 6G). DPM by itself, once again showed enhancing effects on the four metrics, beginning generally after day 16, compared to C9 MN (DMSO). UK-5099 produced little or negative effects on the metrics at early days after addition but provided improvement in the metrics after day 18 (see Discussion). Most importantly, when both compounds were added together, the trajectory of all four metrics followed that of UK-5099 rather than DPM or an intermediate profile, indicating that UK-5099 is dominant, or epistatic, to DPM in its functional effects (Fig. 6G). Given direct interaction and inhibition of MPC by UK-5099, these data provide additional support for the conclusion that the enhancing effects of DPM require MPC function.

## DISCUSSION

Mitochondria (MT) form a dynamic cellular system that is essential for neuronal health; disruption of this system compromises neuronal function and survival and ultimately affects the organism. MT influence cell health in a variety of ways. They regulate intracellular calcium signaling, lipid biosynthesis for cell membranes, the cell cycle, stress responses, innate immunity, nucleotide biosynthesis, amino acid biosynthesis, transcription and translation, catabolic and anabolic metabolism, bioenergetics, and cell death (*69*). Some of their functions are mediated by intrinsic activities and others through physical contact with other organelles and particles, including the endoplasmic reticulum, nucleus, lysosomes, peroxisomes, lipid droplets, Golgi apparatus, mitochondrial derived vesicles, and the plasma membrane (*69*). Given this diversity of roles, therapeutic strategies may benefit from identifying compounds that offset broad MT-system damage and provide a holistic benefit, rather than targeting isolated deficits such as reduced bioenergetics or mitochondrial fragmentation. This systems-level perspective motivated our strategy to identify compounds capable of restoring overall MT-system performance rather than correcting individual mitochondrial defects.

To identify cpds with such “systems-level” effects, we designed a two-tiered assay scheme to audit fundamental functions of the MT-system that are impaired in neurodegenerative diseases. The first employs a multiplexed and longitudinal phenotypic assay using iPSC-derived neurons for MT dynamics, measuring average MT length in the images collected as a surrogate for the balance between fission and fusion, and average MT count as a surrogate for the balance between biogenesis and mitophagy. While MT CA generally correlates with MT count, certain compounds may selectively influence the balance between fission and fusion, altering MT count without changing MT CA. Since neurodegenerative diseases cause the fragmentation of axons and dendrites, we incorporated a fluorescent marker to highlight the neurites, in addition to the one used to highlight MT. Moreover, we monitored neuron viability and death visually, and from the correlated floor level of MT or neurite metrics (Fig. 1). Thus, the first level multiplexed assay produced five metrics describing mitochondrial system integrity: MT count, MT CA, MT length, neurite complexity, and neuron death. Collecting images across days as the neurons differentiate allowed us to follow the progression of pathology and intervene with cpd administration to prevent or arrest it (Fig. 1 and 2). The latter is important because ALS patients typically present after pathology has already begun. Patient-derived iPSC neurons have proven valuable for uncovering MT dysfunction and cellular vulnerabilities in neurodegenerative diseases, supporting iPSC use to study MT system-level effects of candidate cpds (*70–73*). The second level assay measured MT bioenergetics, using the Seahorse MT Stress Test. Measuring ATP generation or the MT IMM potential are alternatives for this (*47, 48*), but do not provide the same depth of information as the MT Stress Test. Although not all MT functions can be assayed given their complexity, these measurements provide a useful overview of MT-system integrity and function. If positive compounds identified from this scheme translate to similar in vivo effects in humans, they will protect from, or arrest, the pathology in MT dynamics and bioenergetics, degeneration of axons and dendrites, MN death, and early death.

Our strategy embraces the “mitochondrial cascade hypothesis” for neurodegeneration, first proposed by Swerdlow and colleagues for sporadic AD (*19, 20*). The hypothesis states that age-related deterioration of MT function, determined by both genetic makeup and environmental factors, influences the probability of AD-related pathology occurring in late-onset AD. Familial AD may be caused by toxic effects of Aβ peptides on MT, or from a sensitized neuronal state due to amyloidosis. MT dysfunction may therefore initiate a cascade of downstream pathological consequences, including axon/dendrite degeneration and neuron death, as observed with diseased iPSC-derived neurons. Other types of molecular/cellular pathology may also arise from the impaired MT-system.

With this perspective and strategy, we identified DPM as a drug that has extraordinary protective and arrestive effects on MT fragmentation and loss of MT content, axon/dendrite degeneration, and the death of iPSC-derived neurons representing two different forms of fALS and one form of fAD. Remarkably, its neuroprotective effects persist for more than 100 days in C9 ALS iPSC-derived MN, whereas untreated MN die at day18/19 (movie S1)! This persistence may partly reflect the stability of DPM in culture media (fig. S5, C and D), although additional factors may contribute. Importantly, the efficacy of DPM extends across neurons representing distinct neurodegenerative disorders. This cross-form efficacy makes sense since MT dysfunction appears to be a universal, and early occurring, hallmark cellular pathology that can account for other types of molecular and cellular pathologies in different neurodegenerative diseases. The cross disease-form efficacy observed with iPSC-derived neurons provides optimism that DPM or related compounds might provide therapeutic value in humans with multiple neurodegenerative diseases. However, the efficacy was far less in AD cortical neurons, suggesting that optimized analogs may need to be designed in a disease-specific manner.

We benchmarked DPM against the two FDA-approved drugs, riluzole and edaravone used broadly for ALS treatment. Riluzole, is an NMDA glutamate receptor blocker, and edaravone, an ROS scavenger. Their benefits are limited: riluzole extends life expectancy by only 2–3 months, and edaravone may slow disease progression with limited improvements in survival and quality of life. Neither drug exhibited pronounced efficacy in our multiplexed assay (fig. S7). Thus, although currently used clinically, neither drug is expected to significantly offset the MT and neurite pathology in patients based on this MN culture model, which may explain their very modest effects in ALS patients (*74–76*).

DPM itself is a very safe drug, currently used as a blood thinner after cardiac surgery and for secondary stroke prevention. There is strong literature support for DPM as an inhibitor of PDEs and ENTs (*55–58*). The anti-clotting effects have been attributed to the combined effects of PDE and ENT inhibition. We tested 12 PDE inhibitors and 3 ENT inhibitors in our MT-focused assays, anticipating that the inhibitors might have similar effects as DPM, thus pointing to a clear mechanism of action and molecular target. However, these assays were negative, except for Zaprinast (fig. S8 and tables S3 and S4), indicating that DPM’s effects are independent of PDE or ENT inhibition. Nor did we obtain evidence that SREBP inhibitors exhibit efficacy in these assays (fig. S8 and table S5).

The observed efficacy of Zaprinast (fig. S9), a known MPC inhibitor, led us to investigate MPC as a potential target for DPM. Docking studies supported possible binding pockets for DPM on MPC, but surprisingly, subsequent functional experiments show that DPM augments, rather than inhibits, pyruvate uptake, perhaps by binding to an allosteric site that stabilizes the IMS-open configuration without preventing pyruvate transport (Fig. 6). Such augmentation would increase the concentration of the substrate pyruvate in the MT, which represents a crucial metabolic nexus. Enzymes that metabolize pyruvate are exclusively localized in the MT matrix where pyruvate is oxidized to acetyl-CoA by pyruvate dehydrogenase to boost oxidative phosphorylation or carboxylated via pyruvate carboxylase to replenish TCA cycle intermediates and precursors for neurotransmitters. Thus, DPM’s ability to enhance MT pyruvate uptake reveals a novel mechanism with potential relevance for neurodegeneration and other diseases associated with MT dysfunction.

The ∼20% increase in pyruvate availability to MT made possible by DPM may appear too small to account for the molecule’s large and dramatic effects on MT and neuronal health observed in the biological assays. However, biological assays reflect a relatively chronic availability of the molecule whereas the biochemical experiments are time limited. Moreover, biochemical effects need not scale proportionally with complex biological outcomes. Interestingly, there are precedents for enhancements in neuronal health by increasing MT pyruvate metabolism. For instance, increasing MT oxidation of pyruvate to acetyl-CoA reduces neuronal cell death and infarction volume after hypoxia ischemic injury (*77*). Nevertheless, there is a balance to be met, since inhibition of MPC by Zaprinast, UK-5099 (above), and other MPC inhibitors can have positive effects on diseased neurons by rewiring MT metabolism to use other energy substrates (*78–80*).

More broadly, these findings suggest that modulation of MT pyruvate transport may represent an underexplored strategy for improving neuronal resilience. MT metabolism lies at the intersection of bioenergetics, biosynthesis, and cellular signaling, and modest changes in substrate availability can propagate through multiple metabolic pathways. By enhancing MT pyruvate uptake, DPM appears to influence MT-system performance at several levels, including bioenergetics, mitochondrial dynamics, and neuronal survival.

The efficacy of DPM in iPSC-derived neurons from ALS and AD patients highlights its therapeutic potential and raises the exciting possibility of drug repurposing (*81, 82*). A major concern about such repurposing is whether DPM crosses the blood brain barrier (BBB) and the blood spinal cord barrier (BSCB). The current evidence for this is conflicting. The earliest studies of DPM pharmacokinetics and central nervous system (CNS) permeability claimed that DPM does not cross the BBB (*83*). This claim was based on the old technique of fluorescence spectrum measurements of DPM in tissue homogenates and has been propagated on drug and chemistry websites and in many subsequent literature reviews. However, DPM has been used successfully in clinical trials for the CNS disorders of restless legs syndrome and schizophrenia (*84–87*). And, it has reported efficacy in animal models for several CNS diseases (*88–94*). Furthermore, direct measurements using sensitive and modern techniques confirm brain penetration in mice in a study on the breast cancer resistance protein (BCRP) in BBB function (*95*). Moreover, the BBB in ALS and AD is leaky (*96–101*), increasing the probability that DPM might exhibit efficacy in patients with these neurodegenerative diseases.

Several considerations should be kept in mind when interpreting these findings. Our experiments rely primarily on iPSC-derived neurons, which represent the human neurons that become dysfunctional and die in neurodegenerative diseases. However, these neurons are cultured in the absence of other neuronal and non-neuronal cell types that make up the central nervous system and other organ systems. In addition, our data support an MPC-dependent mechanism that enhances MT pyruvate uptake, while the precise molecular interaction between DPM and MPC remains to be further defined. Future structural analyses will help refine the understanding of this mechanism, and studies evaluating in vivo functional effects of DPM in ALS models are ongoing to further assess its therapeutic potential.

Taken together, these findings position DPM as a safe, well-characterized drug with a newly recognized MT mechanism that may have therapeutic relevance for neurodegeneration. If BBB penetration proves limiting, DPM may nevertheless provide a valuable scaffold for designing analogs with improved CNS exposure. These observations highlight mitochondrial pyruvate transport as a promising direction for therapeutic development in ALS, AD, and related disorders.

## MATERIALS AND METHODS

### Reagents and Resources

The reagents and resources used for this study are listed in table S6.

### Human iPSCs

The iPSC lines used in this study are described in table S1. All C9 ALS and ISO iPSC lines were cultured in mTeSR1 medium on plates coated with Matrigel (Growth Factor Reduced, 0.083 mg/mL in DMEM/F-12 media with GlutaMAX supplement) at 37°C, 5% CO_2_ and passaged at 60–80% confluency using 0.5 mM EDTA in 1x Dulbecco’s Phosphate Buffered Saline (DPBS without Ca2^+^/Mg2^+^), or ReLeSR or StemPro Accutase (accutase). TDP43^M337V^ HET and REV iPSCs were cultured in StemFlex media on plates coated with 50 µg/mL Synthemax II-SC. All C9 and ISO, TDP43^M337V^ HET and REV iPSC lines were transitioned to mTeSR Plus media and Matrigel. Familial AD patient-specific iPSCs, AG67 (PSEN1^A246E^) and the corresponding isogenic control ISO-AG67 iPSCs were cultured, transduced and differentiated to cortical neurons as previously described (*52*).

### hNIL transfection of human iPSCs for MN differentiation

All C9 and ISO, TDP43^M337V^ HET and REV iPSCs were transfected with the inducible transcription factor transgene cassette (CLYBL-TO-hNIL-BSD-mApple from Michael Ward, Addgene plasmid # 124230; http://n2t.net/addgene:124230) that includes the human transcription factors hNIL (NGN2, ISL1, LHX3) necessary for differentiation of iPSCs into lower motor neurons (MN) (*102*). The transcription factor genes are located behind the tetracycline response element (TRE3G) that is inducible with doxycycline. The vector also contained a CAG promoter driving the constitutive expression of the reverse tetracycline transactivator (rtTA3C) and an EF-1α promoter driving constitutive expression of selection genes (floxed Blasticidin (BSD)-NLS-mApple selection cassette). The vector was inserted at the CLYBL safe harbor locus (*50*).

The iPSCs were grown to 60-80% confluence and then dissociated with accutase into single cells. 5 x10^5^ cells were plated on Matrigel-coated 12-well plates in mTeSR plus media supplemented with 10 μM Y-27632. Media on the cells was changed 2 hours prior to transfection. To prepare the sgRNA-Sp.HiFi Cas9 ribonucleoprotein (RNP) complex, 1 µM of sgRNA (custom sgRNA targeting the human CLYBL intragenic safe harbor locus (between exons 2 and 3) from Synthego (sequence ATGTTGGAAGGATGAGGAAA) was mixed with 1 µM of HiFi SpCas9 protein [Alt-R S.p. HiFi Cas9 Nuclease V3 (Integrated DNA Technologies #1081060)] in Optimem I media and incubated for 5 min at room temperature. iPSCs were transfected with CLYBL-TO-hNIL-BSD-mApple (1 µg/well) and pCE-mp53DD (*103*) (0.2 µg/well, dominant-negative p53, Addgene Plasmid#41856) using Lipofectamine Stem transfection reagent (fig. S2A). Dominant-negative p53 plasmid (episomal) dramatically improves survival of iPSCs that undergo Cas9 dsDNA breaks. The day following transfection, cells were visualized for mApple expression, and if confluent, single cell passaged with accutase to Matrigel-coated 6-well plates for expanding the transfected cells and plated for Blasticidin (BSD) selection (100 -150 µg/mL) in mTeSR plus media supplemented with 10 μM Y-27632. Three days after BSD selection, transfected iPSCs were dissociated with accutase and replated sparsely (1 x 10^4^ cells) onto Matrigel-coated 10-cm dishes in mTeSR plus media supplemented with 10 μM Y-27632 with 100 µg/mL BSD. Individual iPSC clones (mApple positive red fluorescent colonies, fig. S2B) were manually picked under sterile conditions using an EVOS XL Core Imaging System (ThermoFisher Scientific) and transferred to individual wells of a 24-well plate without BSD. All iPSC clones were then expanded in 6-well plates and cryopreserved in CryoStor freezing media.

Each resulting clonal cell line was analyzed for incorporation of the hNIL transgene in the CLYBL locus by PCR (fig. S2C). DNA was extracted from the cells using QuickExtract DNA Extract Solution (Biosearch Technologies, QE09050). PCRs were performed with primers spanning the 5′ and 3′ junctions of the integration, with one primer annealing within the construct and the other outside of the corresponding homology arm. PCR amplifying the intact safe harbor locus was performed to determine whether the clone harbored a heterozygous or homozygous insertion. PCR primer sequences are listed in table S6. Clones with the integration of CLYBL-TO-hNIL-BSD-mApple construct were used for neuronal differentiation. Cell pellets (3 million iPSC per pellet) of the selected iPSC clones were sent for Scorecard, Karyostat+ and Cell ID analysis (ThermoFisher Scientific) to assess cell quality and the presence of abnormalities. Selected C9 iPSC cell pellets were also sent for C9orf72 repeat expansion testing (UCSF Clinical Laboratories).

### Motor neuron differentiation from human iPSCs

All the iPSC clones with stably integrated doxycycline-inducible hNIL transgenes at safe-harbor CLYBL locus were differentiated into lower motor neurons (MN) as described in Fernandopulle et al., (*51*) with modifications (fig. S2). In brief, BSD-selected doxycycline-inducible CLYBL-hNIL iPSCs were dissociated with accutase into single cells and plated in Matrigel-coated plates (for 6-well plate: seed 5–5.5x10^5^ cells/well; for 10-cm dishes: seed 3x10^6^; for 15-cm dishes: 5–7x10^6^) in mTeSR plus medium containing 10 μM Y-27632 (plating day is day –1). Twenty-four hours later (day 0), cells were induced with Neural Induction Medium (NIM): DMEM/F12 with HEPES and glutamine (gln), B27 supplement (1x), MEM Non-Essential Amino Acids (1x, NEAA), GlutaMAX (1x), compound E (200 nM), Y-27632 (10 μM), doxycycline (2 μg/mL) and Culture One (1x). On day 1 of MN differentiation, media was removed and replaced with fresh NIM media containing 2 µg/mL doxycycline. On day 2 of MN differentiation, the cells began showing signs of MN differentiation by beginning to extend neurites. These MN precursors (D2 MNPs) were dissociated with accutase to single cells, strained through 40 µm cell strainer and frozen in Bambanker freezing media (3x10^6^ D2 MNPs/cryovial aliquot).

### iPSC-derived motor neuron culture

Day 2 MNPs were plated on Poly-D-lysine (PDL)/laminin coated 384-well plates along with lentivirus expressing MT-targeted GFP to visualize MT, and a cytoplasmically-targeted mScarlet to visualize somas and neurites. The pCAG-mtTagGFP2-2A-mScarlet plasmid was packaged into a lentivirus by SignaGen Laboratories. PDL-precoated black/clear 384-well plates with 188 µm film bottom and evaporation barrier (Aurora Microplates) were coated with 15 µg/mL laminin diluted in cold Neurobasal Plus media with B27 Plus supplement (1x), NEAA (1×), GlutaMAX (1x), Penicillin/Streptomycin 5000 U (1x), for 2 hrs in incubator at 37°C with 5% CO_2_. D2 MNPs were thawed and replated (10,500 – 14000 D2 C9 and ISO MNPs/well or D2 TDP43^M337V^ REV and HET MNPs/well) in 384-well plates, volume 40 µL) with CAG-mtTagGFP2-2A-mScarlet lentivirus at a MOI=2.5-4 in MN differentiation medium (MNDM): half DMEM/F12 with HEPES and Gln and half Neurobasal Plus neuronal medium, B27 Plus (1x), NEAA (1×), GlutaMAX (1x), Penicillin/Streptomycin 5000 U (1x), compound E (200 nM), doxycycline (dox, 2 μg/mL), laminin (1 μg/mL), BDNF (10 ng/mL), NT3 (10 ng/mL), and GDNF (10 ng/mL) and Culture One (1x). On day 4, 20 µL of the media was removed and an additional 60 µL of fresh MN culture medium was added bringing the total final volume to 80 µL. MN culture medium contained Neurobasal Plus neuronal medium, B27 Plus supplement (1x), NEAA (1x), GlutaMAX (1x), Penicillin/Streptomycin 5000 U (1x), laminin (1 μg/mL), BDNF (10 ng/mL), NT3 (10 ng/mL), and GDNF (10 ng/mL). Every 4 days half of cell culture medium was removed and replaced by fresh MN culture media until day 12. Doxycycline (2 μg/mL) was added only for the day 4 media change and Culture one (1x) for day 4 and/or day 8 media change.

### Immunocytochemistry

The D2 ISO MNPs were thawed and replated in PDL (0.1 mg/mL)/laminin (15 µg/mL) coated 96-well plates at a cell density of 7000 MNPs in 80 μL of MNDM media containing 2 µg/mL dox/well. On day 4, 60 µL of media was removed per well, and 80 µL of fresh MN culture media with 2 µg/mL dox was added (total volume 100 µL). Every 4 days half of cell culture medium was removed and replaced by fresh MN culture media. On day 20, MNs were fixed by adding 4% paraformaldehyde (final concentration) directly to culture media for 15 min at room temperature (RT). MNs were washed twice with DPBS (half volume washes) and permeabilized with 0.5% Triton X-100 in DPBS for 15 min at RT. MNs were washed twice with DPBS and then blocked in 3% bovine serum albumin (BSA) in DPBS for 1 hr at RT and then incubated overnight at 4°C with the primary antibodies described in table S6. The MNs were washed twice with DPBS and incubated with species-specific secondary antibodies at a 1:1000 dilution in BSA blocking buffer and incubated for 1 hr at RT in dark. Cells were washed twice with DPBS and were imaged using 20X objective in IN Cell Analyzer 6000 (GE Healthcare).

### Longitudinal mitochondrial (MT) dynamics assay

D2 MNPs after plating in 384w microtiter plates were imaged every two days, beginning at day 8, and generally until day 30 to follow the MT and neurite metrics across time. Images (four fields per well in 384w plate) were captured using the Image Xpress Micro Confocal High-Content Imaging System (at 37°C, 5% CO_2_) with 60X objective, 0.95 NA with 60 µm pinhole. Each field was acquired as a z-stack of five individual sections of 0.7 µm and maximum intensity projections of each individual stack was used for analysis. A custom segmentation algorithm was applied to each image to segment MT and neurites using the MetaXpress software v.6.5.4.532. The following parameters were collected from these images: total MT count, MT cumulative area (CA), median MT length and neurite CA. CA is the number of pixels representing MT/neurites in each field and provides a metric for MT content/neurite complexity and neurite degeneration. Custom python codes were used to extract the MT and neurite data from these field images and four fields per well were averaged to yield an average per well for each parameter. Each well was a technical replicate with the number of technical replicates indicated in the Figure legends. Experimental average data for each parameter were calculated based on genotype and treatment (with or without compounds). The averaged data for each MT and neurite parameter from each of the time-course experiments, with each experiment being a biological replicate, were then averaged together for each time point tested. A two-way repeated measure ANOVA, followed by Tukey’s or Šídák’s multiple comparisons test was performed to calculate the statistical significance between the groups at each time point using GraphPad Prism (version 10.4.1). To monitor disease progression over time, MNs were imaged every two days starting at day 8. This imaging frequency introduces mild stress that accelerates cell death compared to less frequent imaging. We confirmed this by reducing imaging frequency or the number of sessions, which extended neuronal survival. Both isogenic control and disease-model MNs were subjected to identical imaging conditions to ensure comparability.

### Compound addition, time course and dose response (D:R) assays

The D2 MNPs (C9 ALS and ISO; TDP43^M337V^ and REV) were thawed and replated with CAG-mtTagGFP2-2A-mScarlet lentivirus at a MOI=4 in PDL/laminin coated 384-well plates at a cell density of 10,500-14,000 MNPs in 40 μL of MNDM media containing 2 µg/mL dox/well. On day 4, 20 µL of media with lentivirus was removed per well, and 60 µL of fresh MN culture media with 2µg/mL dox was added (total volume 80 µL). The plates were first imaged on days 8, 10 and 12 as described above. On day 12, to test the selected compounds, C9 MNs and corresponding ISO MNs, were treated with Dimethyl sulfoxide (DMSO) alone (0.125% (v/v), 20-40 technical replicates) or compounds dissolved in DMSO at different concentrations (7-11 point D:R assays with 5 technical replicates). For the experiments to evaluate the effect of DPM on arresting the phenotypes, DPM was added once on days 8, 16, 18, and 20 after the imaging. AG67 (PSEN1^A246E^) and ISO-AG67 cortical neurons were plated and imaged as previously described (*52*). DPM was added on day 10 for the TDP43^M337V^ and REV MNs and on day 8 for AG67 and ISO-AG67 cortical neurons. Prior to adding compounds, 40 µL of media was removed per well, followed by addition of 20 µL of fresh neuron culture media and then 20 µL media containing compounds at the appropriate concentrations. Compounds were placed in a source 384-well plate and then diluted in neuron culture media twice, ensuring that the DMSO concentration remained constant at 0.125% in all wells. Compounds were only added once on the indicated days after imaging. Plates were imaged every two days until day 30, as described above, with no further media changes. Three-to-five independent time-course experiments were carried out for the selected compounds in most experiments. For the D:R experiments, robust Z-scores were calculated relative to the in-plate DMSO control [(Parameter data_compound_ – median_DMSO_)/1.4826 x median absolute deviation_DMSO_] and were averaged for each compound across replicate plates. Robust Z-scores were plotted as D:R curves (across days) for measuring the potency (EC_50_) and efficacy (span) of the compounds.

### Motor neuron viability assay

MN viability in response to DPM (20 µM-156 nM), was evaluated using the RealTime-Glo MT Cell Viability Assay (Promega) according to the manufacturer’s protocol. Viable cells with active metabolism reduce the pro-substrate (MT Cell Viability substrate) into a substrate, which diffuses into the culture medium and is used by NanoLuc luciferase to generate a luminescent signal. The signal corresponds to the number of viable cells. Dead cells do not reduce the pro-substrate and therefore produce no signal. Briefly, NanoLuc luciferase enzyme and a cell-permeant pro-substrate, were added directly to the media of the MNs in culture in 384-well plates on day 31 and luminescence measured on day 33 by CLARIOstar microplate reader (BMG Labtech). A one-way ANOVA, followed by Tukey’s multiple comparisons test was performed to calculate the statistical significance between the groups using GraphPad Prism (version 10.4.1).

### DPM stability assay

Cell culture media from the DMSO/DPM-treated ISO MNs and C9 MNs was collected at the indicated time point and immediately frozen at -80°C. Samples were thawed, and 10 µL of media was added to 70 µL of 90% acetonitrile/10% water containing 20 ng/mL carbamazepine as an internal standard. The samples were allowed to incubate on ice for 15 minutes with occasional shaking to denature and precipitate proteins in the assay media. DPM concentrations were measured by LC-MS/MS on a Sciex 6500 mass spectrometer using an MRM method following the mass transition m/z=505.3→430.2. Concentrations were determined based on a standard curve prepared in assay media analyzed with the media samples. DPM measurements were made by the UF Scripps DMPK scientific core.

### MT bioenergetics assay

Seahorse assay plates (96 wells) were coated with 0.1 mg/mL PDL overnight at 37 °C. Plates were washed with sterile water, air-dried and further coated with 15 µg/mL laminin at 37 °C for 2 hours. Day 2 C9 and ISO MNPs or TDP43^M337V^ and REV MNPs were plated in these plates at a density of 30,000 cells/well (for C9 and ISO MNPs) and 40,000 cells/well (for TDP43^M337V^ and REV MNPs) in a total volume of 180 μL of MNDM medium. A total of 15 replicate wells per neuron type were plated for each experiment. Two days after plating, 135 µL media was removed and replaced with an equal volume of fresh MN culture media. 50% media change was performed every 4 days until the day of the assay. On day 14, 1h prior to the assay, the MN culture media was replaced with Seahorse XF DMEM media supplemented with 10 mM glucose, 2 mM glutamine, and 1 mM pyruvate and the plate was incubated at 37°C in a CO_2_ free incubator. The injection ports of a hydrated Seahorse cartridge were filled with 2 µM oligomycin, 4 µM carbonyl cyanide p-(trifluoromethoxy) phenylhydrazone (FCCP), and 1 µM each of rotenone and antimycin A, and the cartridge was calibrated. Following calibration, the cell culture plate was placed in the Seahorse analyzer, and the oxygen consumption rates (OCR) were measured. For each injection, 3 OCR measurements were recorded. The OCR values were normalized to the number of live cells in each well. Live cell numbers were determined by Calcein AM staining. After the final OCR measurement, Calcein AM was added to each well at a final concentration of 10 µM. The assay plate was incubated in a CO_2_ free incubator at 37°C for 30 min and live cell counts were obtained using an InCell 6000 analyzer (GE Healthcare Technologies, Chicago, IL). Four image fields were collected per well, and the live cells were segmented using a customized segmentation protocol and enumerated using InCell Investigator Developer Toolbox (1.9.2). The total number of cells counted in the four fields was used to normalize the OCR data on a per cell basis and presented as pmol/min/1000 neurons. Four test parameters were analyzed: basal respiration, the general resting state respiratory capacity of the neurons; ATP-linked OCR, an estimation of the amount of energy able to be produced by neurons which is measured after the oligomycin injection; maximal respiration, the amount of energy released when challenged with the electron transport chain uncoupler FCCP; and spare respiratory capacity, the difference between maximal and basal respiration values which indicates the neuron’s energy reserve under periods of stress. Average values of the 15 replicate wells were derived for each experiment for all the four MT respiratory parameters. Three-to-five independent experiments were performed, and unpaired two-sided Student’s *t*-test was performed to calculate the statistical significance between the groups using GraphPad Prism (version 10.4.1). AG67 (PSEN1A246E) and ISO-AG67 cortical neurons were plated, and MT bioenergetics assay was carried out as previously described (*52*).

### Dose response of DPM on MT bioenergetics

Day 2 C9 and ISO MNPs or TDP43^M337V^ and REV MNPs were thawed and plated in PDL/laminin coated 96-well Seahorse assay plates. The MNPs were maintained, and media changes were performed following the protocol described above in the MT bioenergetics section. Day 4 AG67 (PSEN1A246E) and ISO-AG67 cortical neurons were thawed and plated as previously described (*52*). To test the effect of DPM, on day 8, 90µL media was replaced with fresh culture media containing DMSO [0.125% (v/v)] or DPM at the indicated concentrations - 19 nM, 39 nM, 78 nM, 156 nM, and 313 nM for C9 and ISO MNPs; 78 nM, 156 nM, 313 nM, 625 nM and 1.25 µM for TDP43^M337V^ and REV MNs; and 0.625 µM, 1.25 µM, 2.5 µM, 5 µM, and 10 µM for AG67 and ISO-AG67 [5-point D:R assay with 6 – 8 technical replicates]. MT functional tests were performed 6 days after DPM addition on day 14 for C9 and ISO MNs or TDP43^M337V^ REV and HET MNs) and 8 days after DPM addition on day 16 (for AG67 and ISO-AG67) using Seahorse XFe96 analyzer. Three-to-five independent D:R experiments were performed. For the line graphs, Two-way ANOVA, followed by Šídák’s multiple comparisons test was performed to calculate the statistical significance between the groups. D:R curves were plotted to measure the potency (EC_50_) and efficacy (span) of DPM using GraphPad Prism (version 10.4.1). Robust Z-scores were calculated relative to the in-plate DMSO control.

### Molecular Modeling

Molecular docking studies were conducted using the Schrödinger Small-Molecule Drug Discovery Suite (*104–107*). High-resolution crystal structures of MPC in the matrix-open (PDB ID: 9MNW) and IMS-open (PDB ID: 9MNY) conformations were retrieved from the Protein Data Bank (*64*). Protein preparation was performed using Schrödinger’s *Protein Preparation Wizard*, which included adding hydrogen atoms, assigning bond orders, optimizing hydrogen-bonding networks, and performing restrained energy minimization with the OPLS4 force field (*108–110*). The ligand DPM, along with the reference compound GW604714, was prepared using *LigPrep* to generate low-energy, three-dimensional conformations with correct ionization states at physiological pH (7.4). Docking grids were generated to encompass both the pyruvate binding pocket and additional predicted druggable sites identified using *SiteMap*. Docking was carried out using the *Glide* extra-precision (XP) mode, and the predicted binding energies (GlideScore) were recorded. Binding poses were visually inspected to evaluate hydrogen bonding, hydrophobic contacts, and overall fit within the binding site.

### Isolation of liver MT, Oroboros oxygen consumption assay, and immunoblotting

In some experiments, 10-week-old male C57BL6/J mice were used for MT isolation. The generation of *Mpc2* floxed mice has been previously described (*111*). To delete MPC2 in hepatocytes, 2 X 10^11^ particles of adeno-associated virus serotype 8 (AAV8) expressing Cre recombinase under the control of the hepatocyte-specific thyroxine binding globulin (TBG) promoter (AAV8-TBG-Cre; Vector Biolabs; VB1724) was administered intraperitoneally to 6-week-old male *Mpc2* floxed mice to create hepatocyte-specific *Mpc2*-/- mice. Littermate floxed mice receiving 2 X 10^11^ particles of AAV8-TBG-eGFP (Vector Biolabs; VB1743) were used as controls. Mice were studied 3 weeks post-injection of AAV8. All animal experiments were approved by the Institutional Animal Care and Use Committee of Washington University in Saint Louis.

Immunoblotting experiments using a portion of each liver collected for MT isolation were performed to ensure strong or complete knockdown of MPC1/MPC2 expression. Protein was extracted by using radioimmunoprecipitation assay (RIPA) buffer (ThermoFisher Scientific) containing protease inhibitor and phosphatase inhibitor cocktail (Sigma) and mechanical disruption using a Tissuelyser (Qiagen). Protein content was then measured by using a BCA assay. Proteins (10 μg) were separated on a SDS-polyacrylamide gel, transferred to PVDF membranes, incubated with appropriate antibodies, and blots were visualized using a fluorescence imaging system (LI-COR). The following primary antibodies were used: MPC1 (Cell Signaling, 14462), MPC2 (Cell Signaling, 46141), and vinculin (Cell Signaling, 13901). The secondary antibody used was IRDye 800CW goat anti-rabbit IgG (LI-COR, 926–32211).

MT isolation and respiration studies were performed as previously described (*112*). In brief, the left lobe of the liver was excised from mice after CO_2_ asphyxiation and homogenized in buffer containing 250 mM sucrose, 10 mM Tris base, and 0.5 mM EDTA (pH 7.4) by 8–10 passes of a glass-on-glass Dounce homogenizer on ice. Homogenates were centrifuged at 1000 x *g* for 5 min at 4°C to pellet nuclei and undisrupted cells. The supernatants were then centrifuged at 10,000 × *g* for 10 min at 4°C to enrich for MT; this MT pellet was washed and re-pelleted twice in fresh sucrose/tris buffer. The MT pellet was then solubilized in 300 μL of Mir05 respiration buffer (0.5 mM EGTA, 3 mM MgCl_2_, 60 mM lactobionic acid, 20 mM taurine, 10 mM KH_2_PO_4_, 20 mM HEPES, 110 mM sucrose and 1 g/L of fatty acid free bovine serum albumin; pH 7.1). MT protein content was then measured by BCA assay, and 50 µg of MT protein was added to the chambers of an Oxygraph O2K (Oroboros Instruments), with a total volume of 2 mL Mir05 buffer set to 37°C. Respiration was stimulated with 5 mM pyruvate/2 mM malate until a steady state was reached and then 2 mM ADP was added. After again reaching steady state respiration rates (baseline), UK-5099 or DPM were added to the chamber at sequentially increasing indicated concentrations. Steady-state rates of oxygen consumption were assessed for 1–2 min before addition of the next highest dose of UK-5099 or DPM. Oxygen consumption rates (OCR) were calculated from the change in oxygen concentration over time and normalized to 50 µg of MT within the chamber.

### Isolation of liver MT and the pyruvate uptake assay

Four-month-old male C57BL6/J mice were used for MT isolation in accordance with UF Scripps Institutional Animal Care and Use Committee. Liver from one mouse was excised and homogenized on ice in MSHE buffer supplemented with 0.5% BSA (MSHE: 210 mM mannitol, 70 mM sucrose, 5 mM HEPES, 1 mM EGTA, pH 7.4) using a pre-chilled glass homogenizer with 3 passes of motor-driven pestle set at 200 rpm. The homogenate was centrifuged at 1,000 × *g* for 5 min at 4°C, filtered through wet cheesecloth, and centrifuged at 11,600 × *g* for 10 min at 4°C. The resulting MT pellet was washed twice in MSHE buffer and centrifuged at 11,600 × *g* for 10 min at 4°C. The purified MT pellet was resuspended in 1.4 mL of MSHE buffer, and protein concentration was determined by BCA assay. The MT protein yield was between 18 - 20 mg/liver.

For the pyruvate uptake assay, modified from Gray et al., (*68*) the MT solution was divided equally into seven microcentrifuge tubes and pelleted at 8,000 × *g* for 15 min at 4°C. Each MT pellet was resuspended in uptake buffer (120 mM KCl, 5 mM HEPES, 1 mM EGTA, 2 µM rotenone, 2 µM antimycin A, pH 7.4) so that the final concentration was 7.7 µg/µL. Both UK-5099 and DPM were tested in dose response. A portion of each resuspended pellet was incubated with the following concentrations of either UK-5099 or DPM: 0 µM/DMSO, 0.313 µM, 0.625 µM, 1.25 µM, 2.5 µM, 5 µM and 10 µM. Another portion of each resuspended pellet was also incubated with 10 µM UK-5099 that served as the negative control reaction. After a 5 min incubation with UK-5099, or a 3-hour incubation with DPM to allow time for DPM to diffuse across the MT membranes, 30 µL of substrate mix (200 µM sodium pyruvate and 0.176 µmol C^14^-pyruvate) in uptake buffer pH 6.1) was added to 30 µL of the MT suspension for a final substrate concentration of 100 µM. The reaction was stopped after 20 sec with 140 µL of stop solution (10 µM UK-5099 in uptake buffer, pH 6.8). Three technical replicates of the test substance (UK-5099 or DPM) and two UK-5099 negative control technical replicates were performed for each test substance concentration. The stopped reactions were centrifuged at 21,000 × *g* for 3 min to pellet the MT. The pellets were washed once with 200 µL of 10 µM UK-5099, 100 µM sodium pyruvate in uptake buffer, pH 6.8, and then resuspended in 500 µL of 1x RIPA buffer. MT lysates were transferred to scintillation vials with 13 mL scintillation fluid and counted. Extra MT in each suspension was assayed again for protein concentration using a BCA assay. Raw C^14^ counts per min were converted to pmol pyruvate/mg protein using the post-assay BCA assay values. The average UK-5099 treated negative control values were subtracted from the average across the technical replicates for each test substance concentration. Three independent experiments were performed for UK-5099 and five independent experiments for DPM. A mixed-model ANOVA was performed using the standard least squares model in JMP v15. Post hoc analysis was performed with Student’s *t*-test.

### Quantification and statistical analysis

Data were analyzed with GraphPad Prism v10.4.1. Data for the MT dynamics assays are presented as box plots with individual data points that are mean of 3-to-5 independent time-course experiments. Data for the MT bioenergetics, and Oroboros oxygen consumption assay are shown as mean ± SEM unless specified otherwise. *p* ≤ 0.05 was defined as statistically significant. For comparison of normally distributed data from two groups, an unpaired two-sided Student’s *t*-test was performed. For the statistical analysis of more than two groups, one-way ANOVA with Tukey’s multiple comparisons test, Two-way ANOVA or Two-way repeated measure ANOVA with Tukey’s multiple comparisons test or Šídák’s multiple comparisons test was performed. Data for the pyruvate uptake assay were analyzed using the mixed-model ANOVA with standard least squares model in JMP v15 with Student’s *t*-test for Post hoc comparisons. *p<0.05, **p<0.01, ***p<0.001, ****p<0.0001; ns, non-significant difference.

## Supporting information

All Supplemental figures and Tables

## Acknowledgements

We thank Christopher Grunseich (NINDS/NIH), and Michael Ward (NINDS/NIH) for providing the CLYBL-TO-hNIL-BSD-mApple plasmid and useful advice on motor neuron differentiation. We would also like to thank Bhuvaneish Selvaraj, University of Edinburgh for sharing the Becker S6, and M211 R2 iPSC lines and Jared L. Sterneckert, University of Dresden for sharing 33.1 iPSC line. We thank Farid Chehab and his team, UCSF Clinical Laboratories, for C9orf72 repeat expansion testing of ALS iPSCs. Statistical consultation was provided by Dr. Gogce Crynen at the UF Scripps Bioinformatics Core. We thank the UF Scripps DMPK scientific core for their assistance in measuring DPM in cell culture media.

## Funding

This work was supported by National Institutes of Health grants R35NS097224 (R.L.D.), R33AG068887 (R.L.D), RF1AG077160-01A1 (B.E.), R01DK104735 (B.N.F.), P30DK056341 (B.N.F.), UF Scripps DMPK scientific core (NIH grant 1 S10OD030332-01), and the Community Foundation for Palm Beach and Martin Counties (R.L.D.).

## Author Contributions

Conceptualization: N.Shahani, B.N.F., R.L.D.

Methodology: N.Shahani., R.B., C.M., N.Sharma., M.H., N.N., H.D.W., P.Z., B.E., L.H., M.D.C., T.D.B., B.N.F., R.L.D.

Project administration: R.L.D.

Supervision: N.Shahani., R.L.D., B.N.F.

Writing – original draft: N.Shahani., R.L.D.

Writing – All authors reviewed and edited the manuscript.

Funding acquisition: R.L.D., B.E., B.N.F., M.D.C

## Competing Interests

B.N.F. is a shareholder and a member of the scientific advisory board of Cirius Therapeutics. B.E. is a cofounder and holds stock in Pelagos Pharmaceuticals, Inc. R.L.D is the founder and a shareholder of Mitochondrial Medicines Corporation. The research presented here is related to US patent applications 17/756,842 and PCT/US26/15790.

## Data, code, and materials availability

All data and code needed to evaluate and reproduce the results in the paper are present in the paper and/or the Supplementary Materials. The generated materials and code used in this study are available by request from Ronald Davis or Neelam Shahani.

