## Supplementary material for "Pharmacological rescue of mitochondrial dysfunction, neurite degeneration, and premature death of ALS and AD iPSC-derived neurons": All Supplemental figures and Tables

Neelam Shahani *et al.*

**This PDF file includes:**

Figs. S1 to S9
Tables S1 to S6
Legend for movie S1

**Other Supplementary Materials for this manuscript include the following:**

Movies S1

**Fig. S1.**

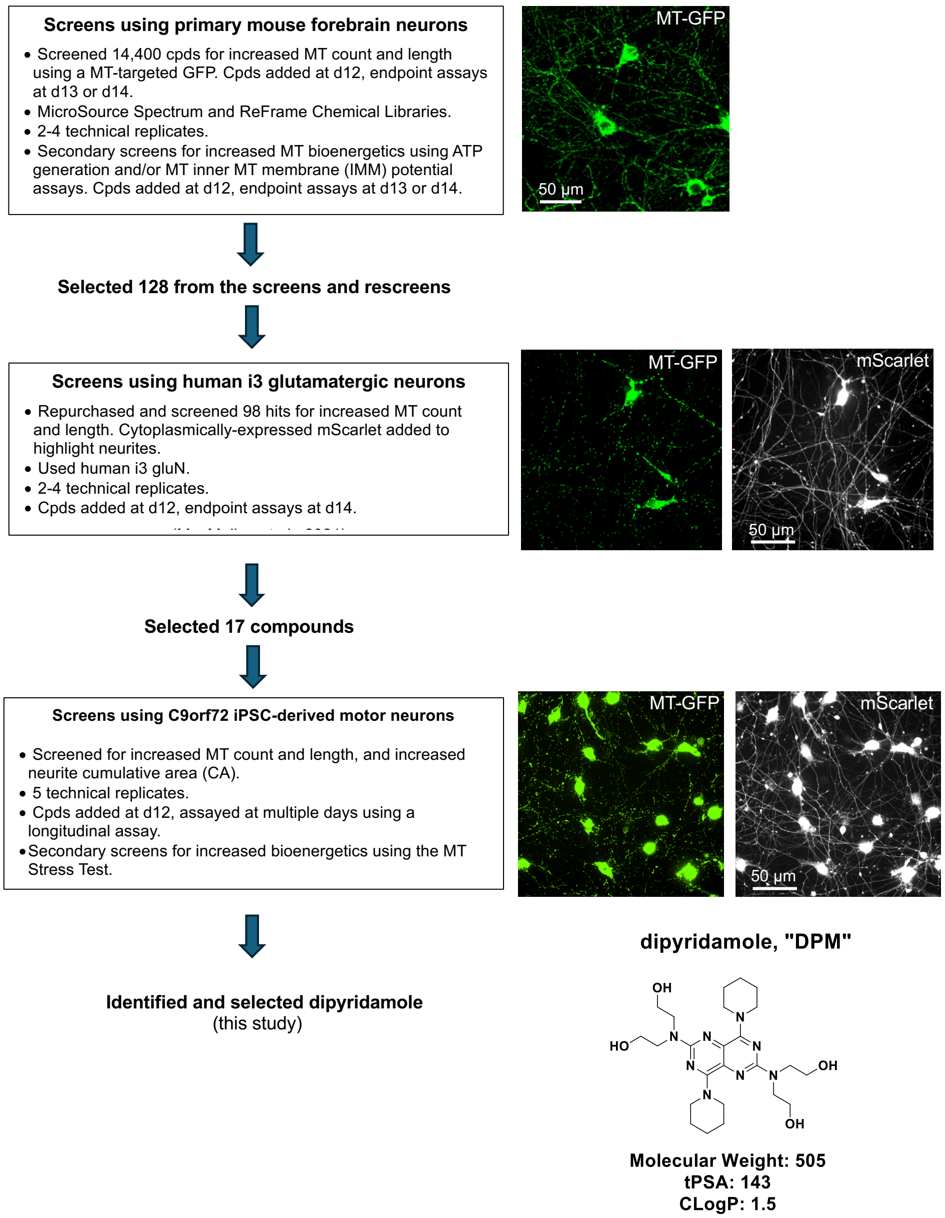

**Fig. S1.** Compound progression and triage.

**Fig. S1. Compound progression and triage**. The selection of compounds (cpds) for broad effects on the neuronal population of MT from the initial screens using mouse forebrain neurons, to human iPSC-derived glutamatergic neurons, to iPSC-derived motor neurons for C9orf72 ALS. Metrics collected include MT length, as a surrogate for the balance between fission and fusion; MT count, as a surrogate for the balance between biogenesis and mitophagy; and in the iPSC-based neuron screens, neurite cumulative area (CA) to monitor the integrity of the axons and dendrites. Dipyridamole (DPM) was a final hit from the progression that is detailed in this report.

**Fig. S2.**

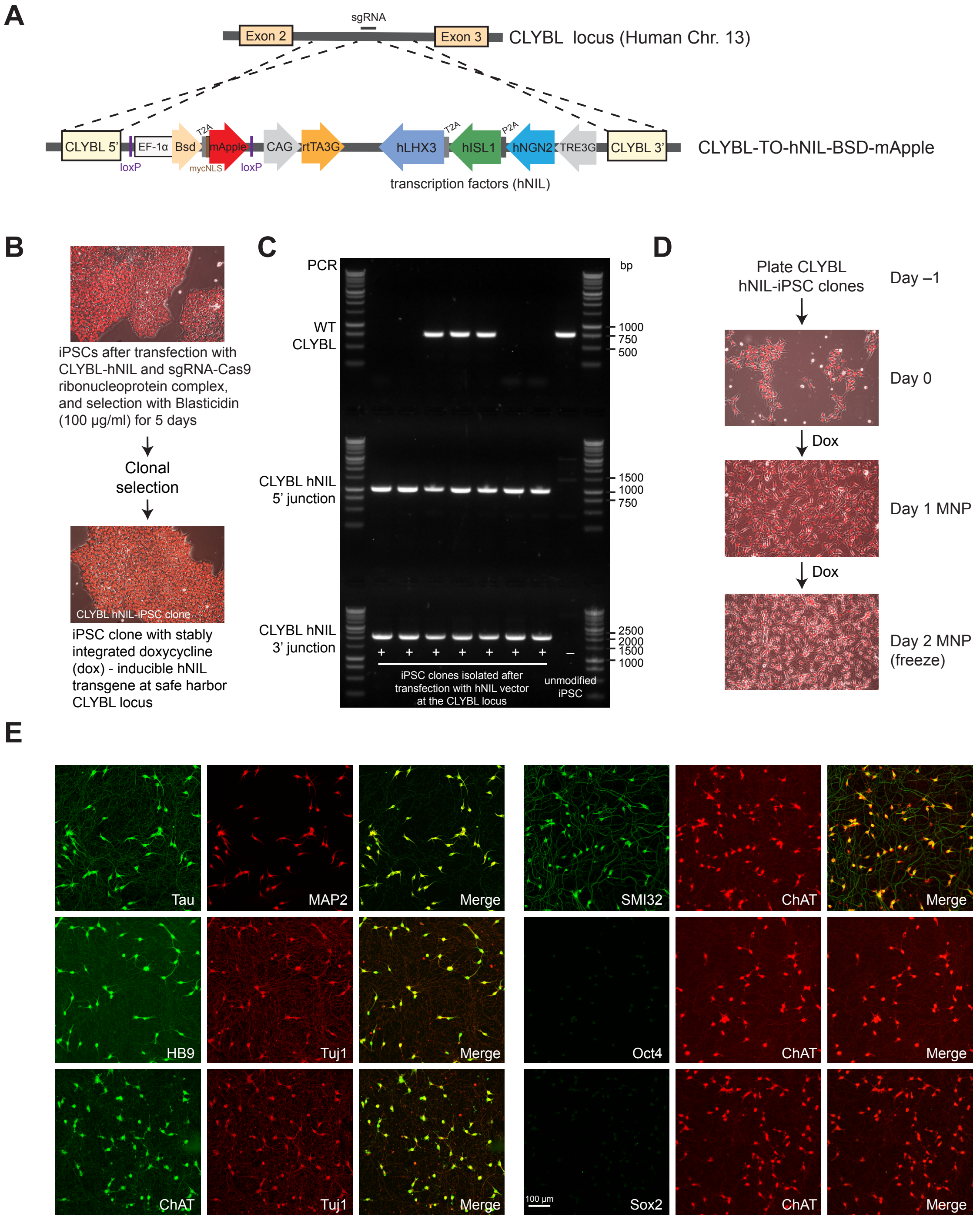

**Fig. S2. Lower motor neuron (MN) differentiation from human iPSCs.**

**Fig. S2. Lower motor neuron (MN) differentiation from human iPSCs**. **(A)** Schematic of the donor construct containing doxycycline (dox)-inducible transcription factors (hNIL: NGN2, ISL1, LHX3) stably integrated into the CLYBL safe-harbor locus using CRISPR/Cas9. **(B)** Representative images for the generation of the iPSC clones with the hNIL transgene at the CLYBL locus (CLYBL hNIL-iPSC clones). **(C)** Confirmation of targeted integration of hNIL cassette at the CLYBL safe harbor locus in iPSC lines, with 3 clones heterozygous for the construct and 4 homozygous (no WT CLYBL band). **(D)** The time course from plating CLYBL hNIL-iPSC clones to induce differentiation into MN progenitors (MNP) in the neural induction media containing 2 µg/ml dox for 2 days. D2 MNPs are frozen in convenient-sized aliquots and later replated on poly-D-lysine (PDL)/laminin-coated plates in MN culture media to generate lower motor neurons (LMNs). **(E)** Representative immunofluorescent images of day 20 isogenic control-iPSC derived MNs stained for pan neuronal (tau, MAP2, Tuj1) and lower MN markers (HB9, ChAT, SMI32). ChAT positive MN do not express pluripotency markers, Oct4 and Sox2. 20X magnification, Scale bar = 100 µm

**Fig. S3.**

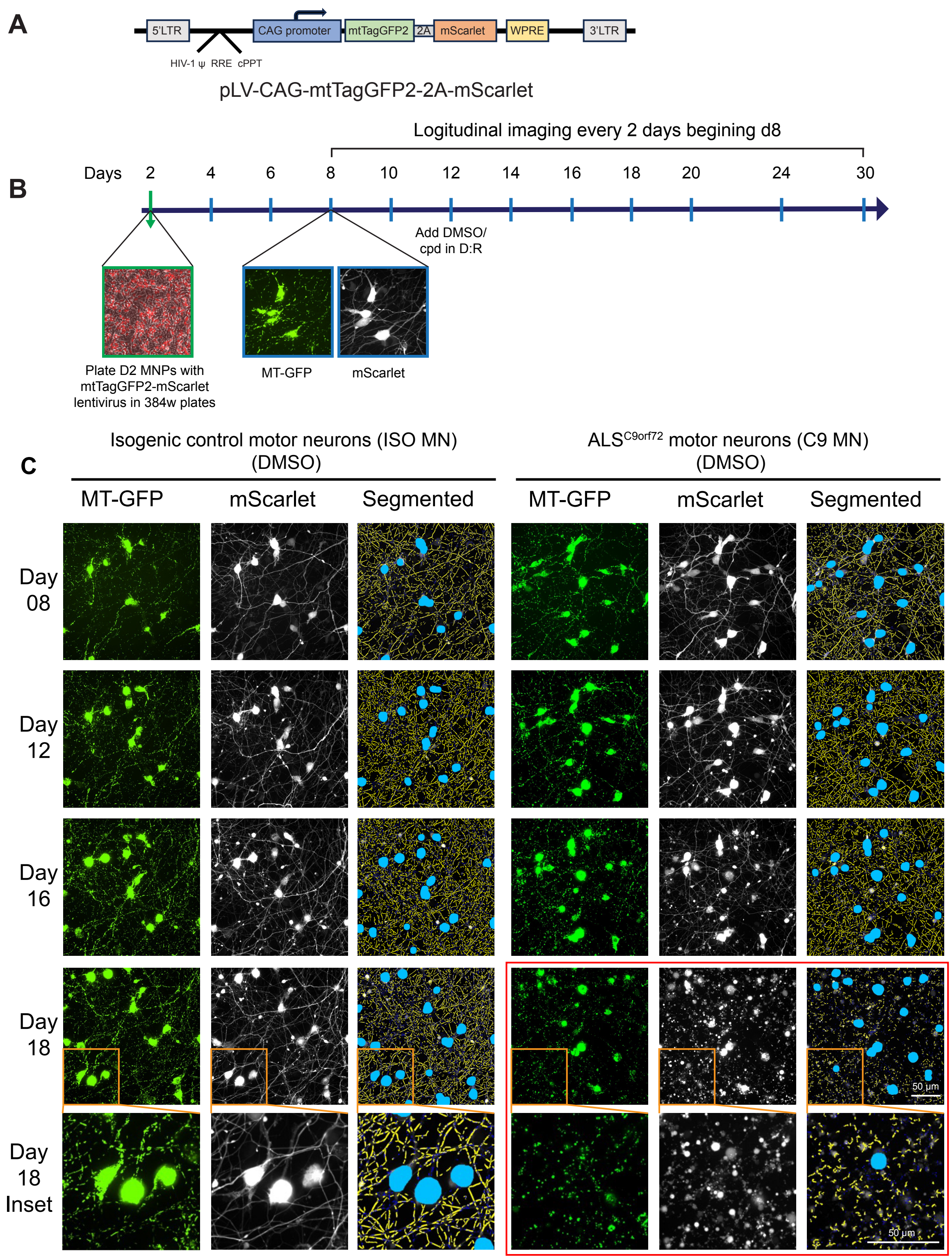

**Fig. S3. MT/neurite dynamics assay overview.**

**Fig. S3. MT/neurite dynamics assay overview. (A)** The pLV-CAG-mtTagGFP2-2A-mScarlet plasmid components that were packaged into a lentiviral vector. **(B)** Overview of the MT and neurite dynamics assay timeline: D2 MNPs were plated in 384-well plates with mtTagGFP2-mScarlet lentivirus at an MOI that provided for sparse labeling, followed by longitudinal imaging every two days starting on day 8. Compound (cpd)/DMSO was added on day 12. **(C)** Images obtained from the same set of DMSO-treated ISO and C9 MN across days of differentiation. Representative images showing the neuronal MT-GFP in green, cytoplasmic mScarlet fluorescence in white, and the images after object segmentation. MN somas (blue in the 3rd and 6th columns) are masked out to focus on neurites. In these columns, the segmented neurites are yellow, and the MT are small dark blue dots on top of the yellow neurites (see day 18 inset). By day 18, there is an obvious loss of MT and neurite content in the C9 MN (images within the red box) when compared to ISO MN. 60X magnification, Scale bar = 50 μm.

**Fig. S4.**

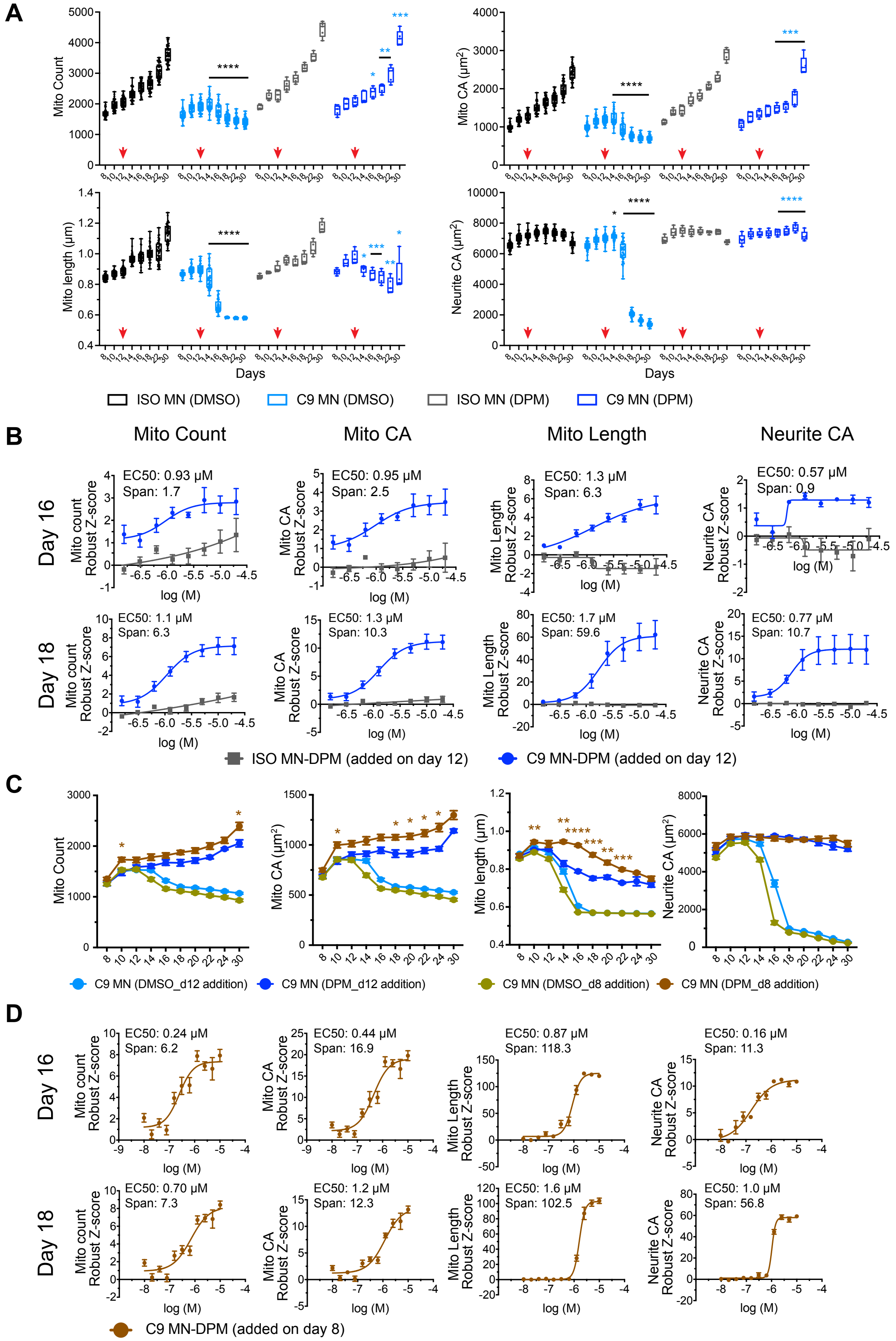

**Fig. S4. DPM is neuroprotective for C9 MN.**

**Fig. S4. DPM is neuroprotective for C9 MN. (A)** Effect of DPM on the MT count, MT CA, MT length and neurite CA in the longitudinal assay using ISO MN and C9 MN. Data are box plots from one time-course experiment with n = 44 technical replicates for DMSO and n = 5 technical replicates for DPM-treated groups. The box extends from the 25th to 75th percentiles, with the line in the box indicating the median. The whiskers extend to the minimum and maximum values. Red arrows indicate the addition of single dose of DMSO (0.125%) / DPM (10 µM) on day 12. *p<0.05, **p<0.01, ***p<0.001, ****p<0.0001, Two-way repeated measures ANOVA, Tukey's multiple comparisons test [black asterisk C9 MN (DMSO) vs ISO MN (DMSO); blue asterisk C9 MN (DPM) vs C9 MN (DMSO)]. **(B)** D:R curves for DPM at day 16 and day 18 using C9 MN and ISO MN. D:R data (mean ± SEM of 5 independent experiments) for MT count, MT CA, MT length, and neurite CA are presented as robust Z-scores using C9 MN / ISO MN relative to the respective in-plate DMSO control wells. DPM was added on day 12 and tested from 156 nM-20,000 nM concentrations. Robust Z-scores were plotted against the $\log_{10}$ of the molar concentrations of DPM. The measured $EC_{50}$ values and spans for the four parameters are shown. Potency is determined by $EC_{50}$, while efficacy is measured by span. Span is the y-axis score between the baseline and the asymptotic value. Robust Z-scores ≥2 or ≤-2 are considered significant. **(C)** Effect of DPM added on day 8 versus day 12. Brown asterisks illustrate the significant differences of the C9 MN (DPM added on d8) vs C9 MN (DPM added on d12). *p<0.05, **p<0.01, ***p<0.001, ****p<0.0001, Two-way repeated measures ANOVA with Tukey's multiple comparisons test [C9 MN (DMSO d12 addition, n = 42, C9 MN (DPM d12 addition, 10 µM, n = 5), C9 MN (DMSO d8 addition, n = 21) and C9 MN (DPM d8 addition, 10 µM, n = 5 technical replicates). **(D)** D:R data (mean ± SEM) for MT count, MT CA, MT length, and neurite CA are presented as robust Z-scores using C9 MN relative to in-plate DMSO control wells. DPM was added on day 8 and tested at concentrations ranging from 10 nM to 10,000 nM. The measured $EC_{50}$ values and spans for the four parameters are shown.

**Fig. S5.**

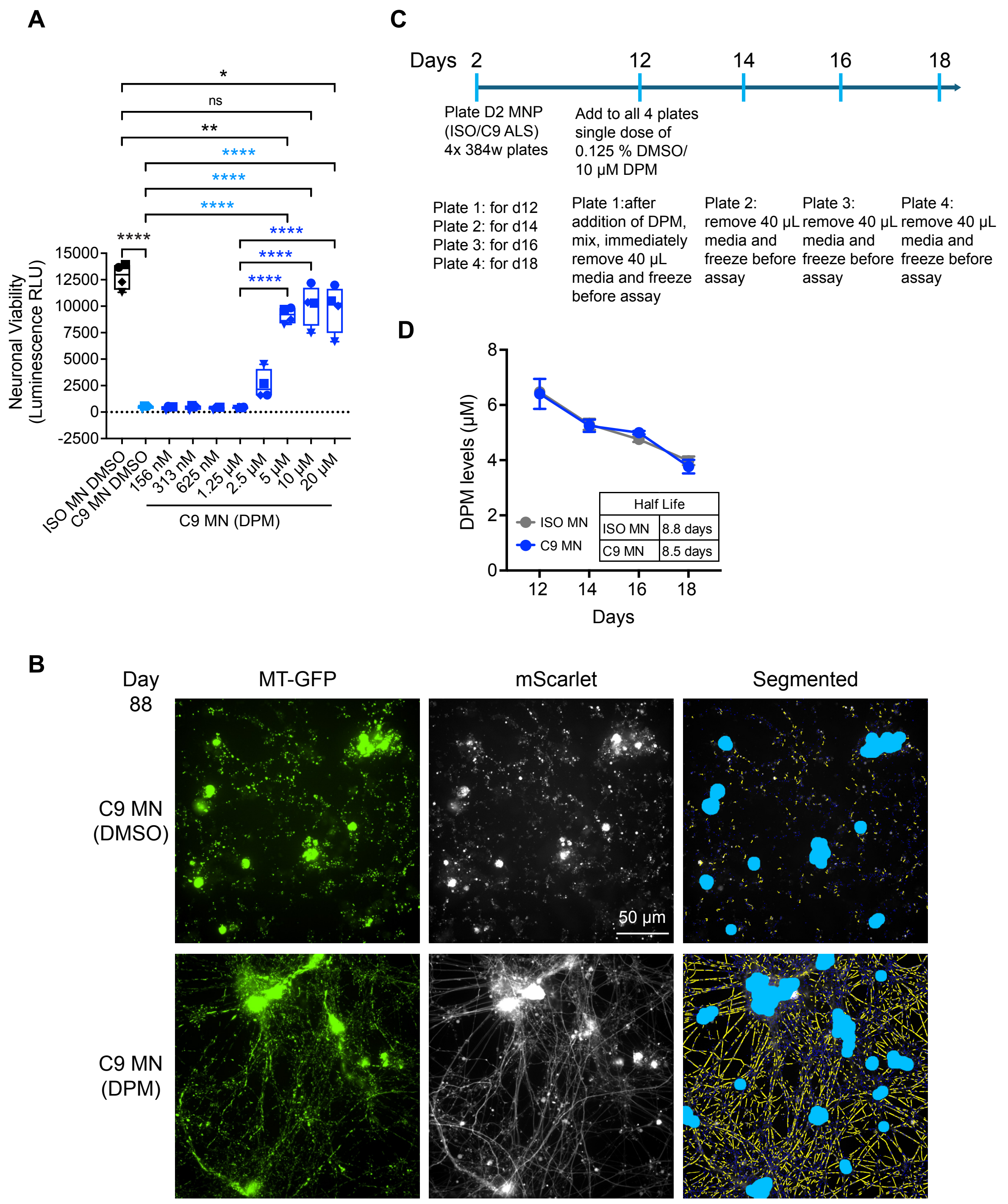

**Fig. S5. DPM increases neuronal survival of C9 MN.**

**Fig. S5: DPM increases neuronal survival of C9 MN. (A)** Dose-dependent effect of DPM on neuronal viability in C9 MN. Data obtained with the RealTime-Glo MT Cell Viability reagent added at day 31 and assayed at day 33. Data are presented as box plots with individual data points (unique symbols) from n = 4 independent experiments. Box plots show the median (center line), the 25th to 75th percentiles (box), and minimum and maximum values (whiskers). Black asterisks illustrate the significant differences of the C9 MN (DMSO) vs ISO MN (DMSO). Light blue asterisks illustrate the significant differences of the C9 MN (DPM) vs ISO MN (DMSO). Dark blue asterisks illustrate the significant differences of the C9 MN (DPM 20 μM, 10 μM, 5 μM) vs C9 MN (DPM 1.25 μM). *$p<0.05$, **$p<0.01$, ****$p<0.0001$, One-way ANOVA with Tukey's multiple comparisons test (n = 4 independent experiments) with n = 16 replicates (DMSO group) and n = 5 replicates (DPM group) in each experiment. ns = not significant. **(B)** Representative images of C9 MNs at day 88, with a single dose of DPM (10 μM) or DMSO (0.125%) added on day 12. **(C)** Schematic of the experiment to test DPM's stability in the ISO MN and C9 MN culture media. **(D)** Quantification of the DPM levels in media at days 12, 14, 16, 18 from ISO and C9 MNs with DPM added on day 12 (as described in A). Data are mean ± SEM, two independent experiments with n = 5 technical replicates per experiment for each group.

#### Fig. S6.

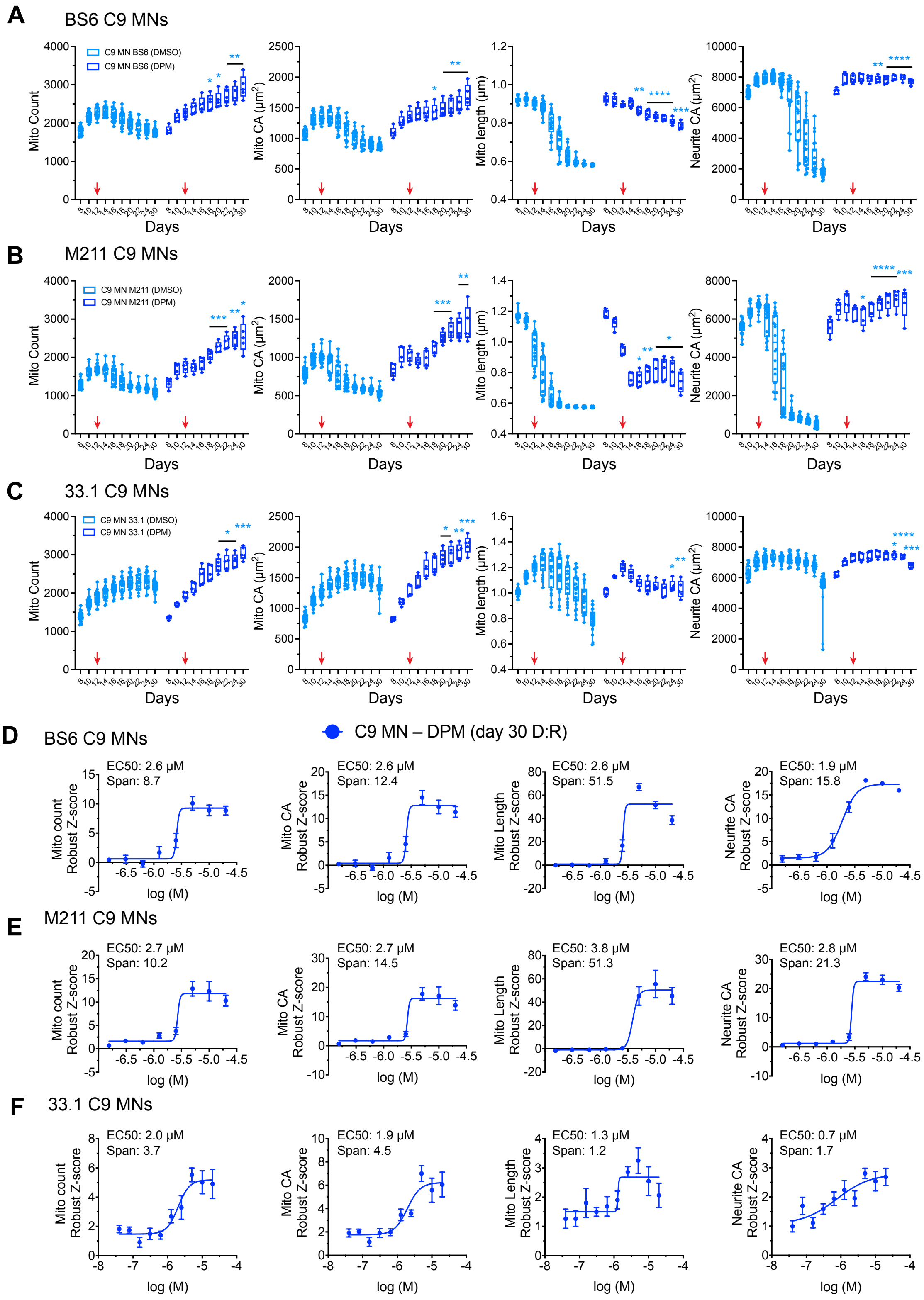

**Fig. S6. DPM is neuroprotective in MNs from three other independent C9orf72 iPSC lines.**

**Fig. S6: DPM is neuroprotective in MNs from three other independent *C9orf72* iPSC lines. (A–C)** Effect of DPM on MT count, MT CA, MT length and neurite CA using BS6 C9 MNs **(A)**, M211 C9 MNs **(B)**, and 33.1 C9 MNs **(C)**. Data are box plots from one time-course experiment showing the median (center line), 25th–75th percentiles (box), and minimum and maximum values (whiskers). Asterisks illustrate the significant differences of the C9 MN (DPM) vs C9 MN (DMSO). *$p<0.05$, **$p<0.01$, ***$p<0.001$, ****$p<0.0001$, Two-way repeated measures ANOVA with Šídák's multiple comparisons test [n = 22 technical replicates (DMSO) and n = 5 technical replicates (DPM)]. **(D–F)** D:R curves for DPM at day 30 using BS6 C9 MNs **(D)**, M211 C9 MNs **(E)**, and 33.1 C9 MNs **(F)**. D:R data (mean ± SEM) for MT count, MT CA, MT length, and neurite CA (robust Z-score using C9 MN relative to in-plate DMSO control wells) using DPM- treated BS6, M211, 33.1 C9 MNs. DPM was added on day 12 and tested at concentrations ranging from 156 nM-20,000 nM. Data are shown only for day 30. The measured $EC_{50}$ values and spans for the four parameters are shown.

**Fig. S7.**

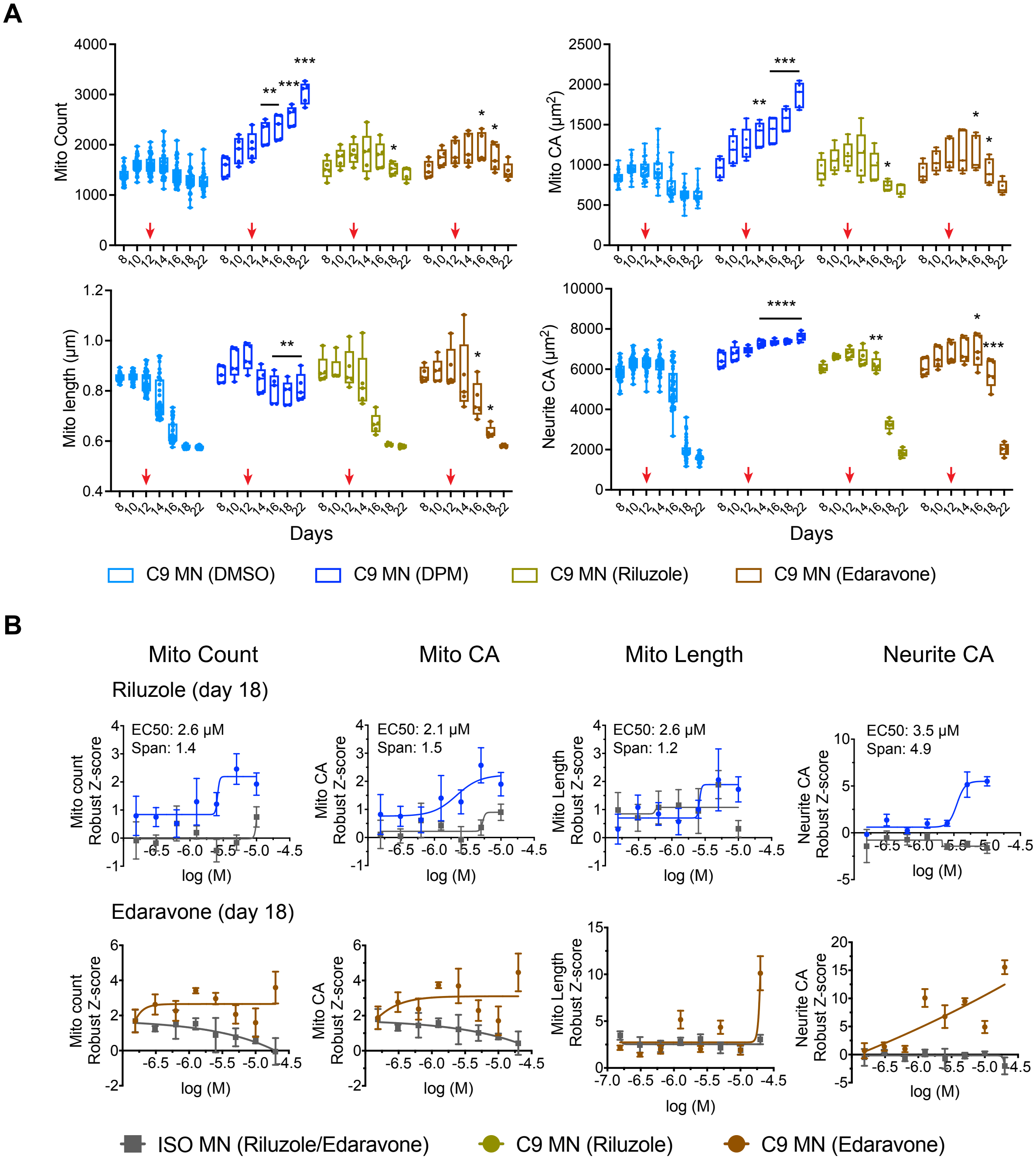

**Fig. S7. Riluzole or Edaravone have very modest effect on the MT/neurite phenotypes in C9 MN.**

**Fig. S7: Riluzole or Edaravone have very modest effects on the MT/neurite phenotypes in C9 MN. (A)** Effect of Riluzole, Edaravone, and DPM added on day 12 on

MT count, MT CA, MT length, and neurite CA in the longitudinal assay using C9 MNs. Data are box plots from one time-course experiment showing the median (center line), 25th–75th percentiles (box), and minimum and maximum values (whiskers). Asterisks illustrate the significant difference of the C9 MN (DPM/Riluzole/Edaravone) vs C9 MN (DMSO). *p<0.05, **p<0.01, ***p<0.001, ****p<0.0001, Two-way repeated measures ANOVA with Šídák's multiple comparisons test, n = 42 replicates (DMSO), n = 5 replicates (DPM/Riluzole/Edaravone). Red arrows indicate the addition of single dose of DMSO (0.125%) / DPM (10 µM) / Riluzole (10 µM) / Edaravone (20 µM) added on day 12. **(B)** D:R data (mean ± SEM) for MT count, MT CA, MT length, and neurite CA (robust Z-score using ISO MN/C9 MN relative to respective in-plate DMSO control wells) using riluzole or edaravone-treated ISO/C9 MN. Riluzole was tested at concentrations ranging from 156 nM to 10,000 nM, while Edaravone was tested at concentrations ranging from 156 nM to 20,000 nM. Data are shown only for day 30. The measured $EC_{50}$ values and spans for the four parameters are shown.

**Fig. S8.**

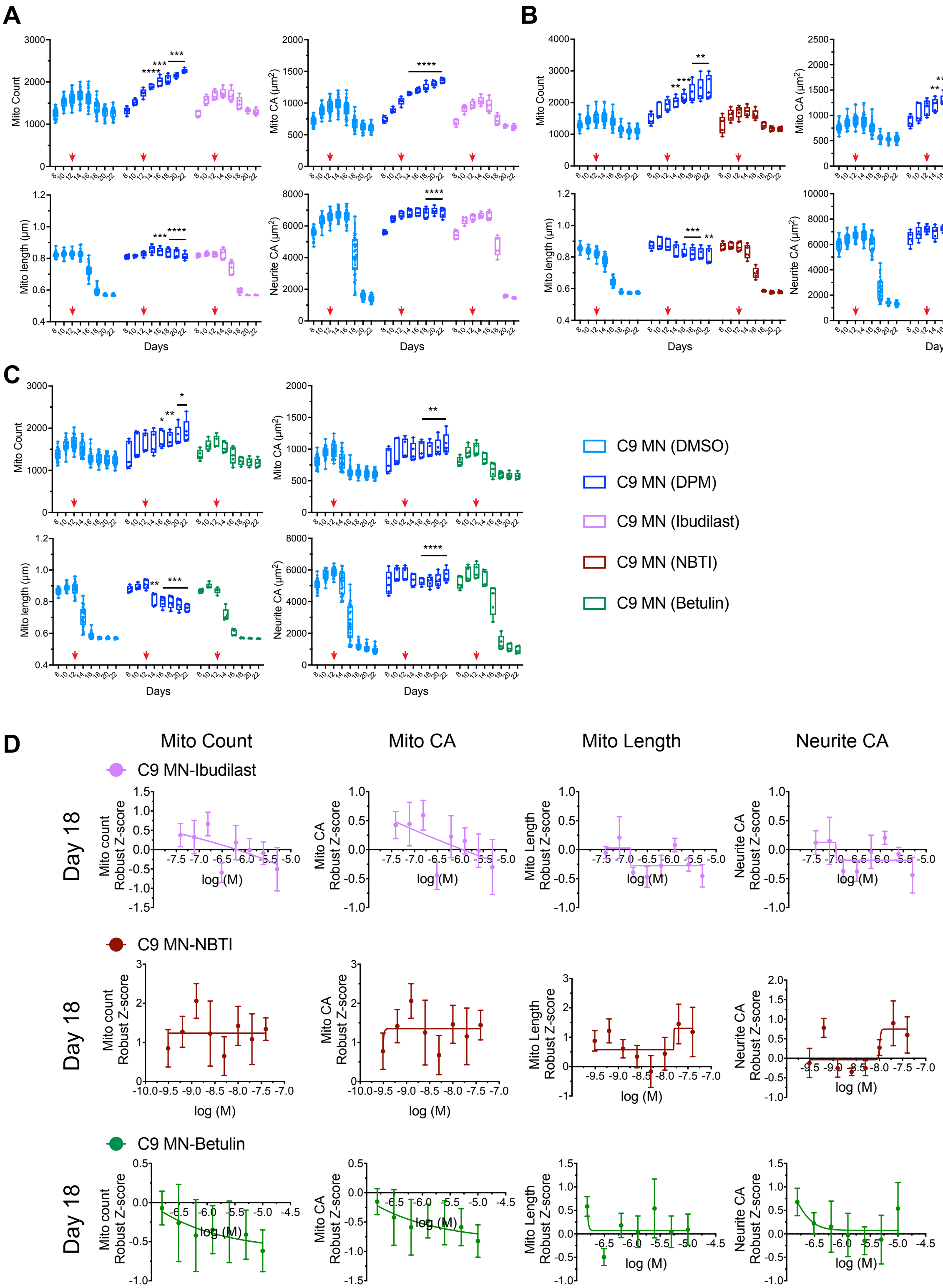

**Fig. S8. Time course and D:R experiments for Ibudilast (PDE inhibitor), NBTI (ENT inhibitor), and Betulin (SREBP inhibitor) in C9 MN.**

**Fig. S8: Time course and D:R experiments for Ibudilast (PDE inhibitor), NBTI (ENT inhibitor), and Betulin (SREBP inhibitor) in C9 MN. (A-C)** Effect of Ibudilast **(A)**, NBTI **(B)**, Betulin **(C)**, and DPM added on day 12 on MT count, MT CA, MT length, and neurite CA in the longitudinal assay using C9 MNs. Data are box plots from one time-course experiment showing the median (center line), 25th–75th percentiles (box), and minimum and maximum values (whiskers). Asterisks illustrate the significant difference of the C9 MN (DPM/Ibudilast/NBTI/Betulin) vs C9 MN (DMSO). ***$p<0.001$, ****$p<0.0001$, Two-way repeated measures ANOVA with Tukey's multiple comparisons test, n = 42-44 replicates (DMSO), n = 5 replicates (DPM/Ibudilast/NBTI/Betulin). Red arrows indicate the addition of single dose of DMSO (0.125%) / DPM (10 µM) / Ibudilast (78 nM)/ NBTI (0.625 nM)/ Betulin (5 µM) added on day 12. **(D)** Ibudilast/NBTI/Betulin D:R data for the MT count, MT CA, MT length and neurite CA at day 18 presented as robust Z-scores using C9 MN relative to in- plate DMSO control wells. Ibudilast, tested at concentrations ranging from 39 nM to 5,000 nM, is an inhibitor of PDE 3AB, 4ABCD, and 5A, with $IC_{50}$ values ranging from 54 nM to 3510 nM. NBTI, tested at concentrations ranging from 0.313 nM to 40 nM, is an ENT1/2 inhibitor with an $IC_{50}$ of 0.4 nM for hENT1. Betulin, tested at concentrations ranging from 156 nM to 10,000 nM is a SREBP inhibitor with an IC50 of 14500 nM.

**Fig. S9.**

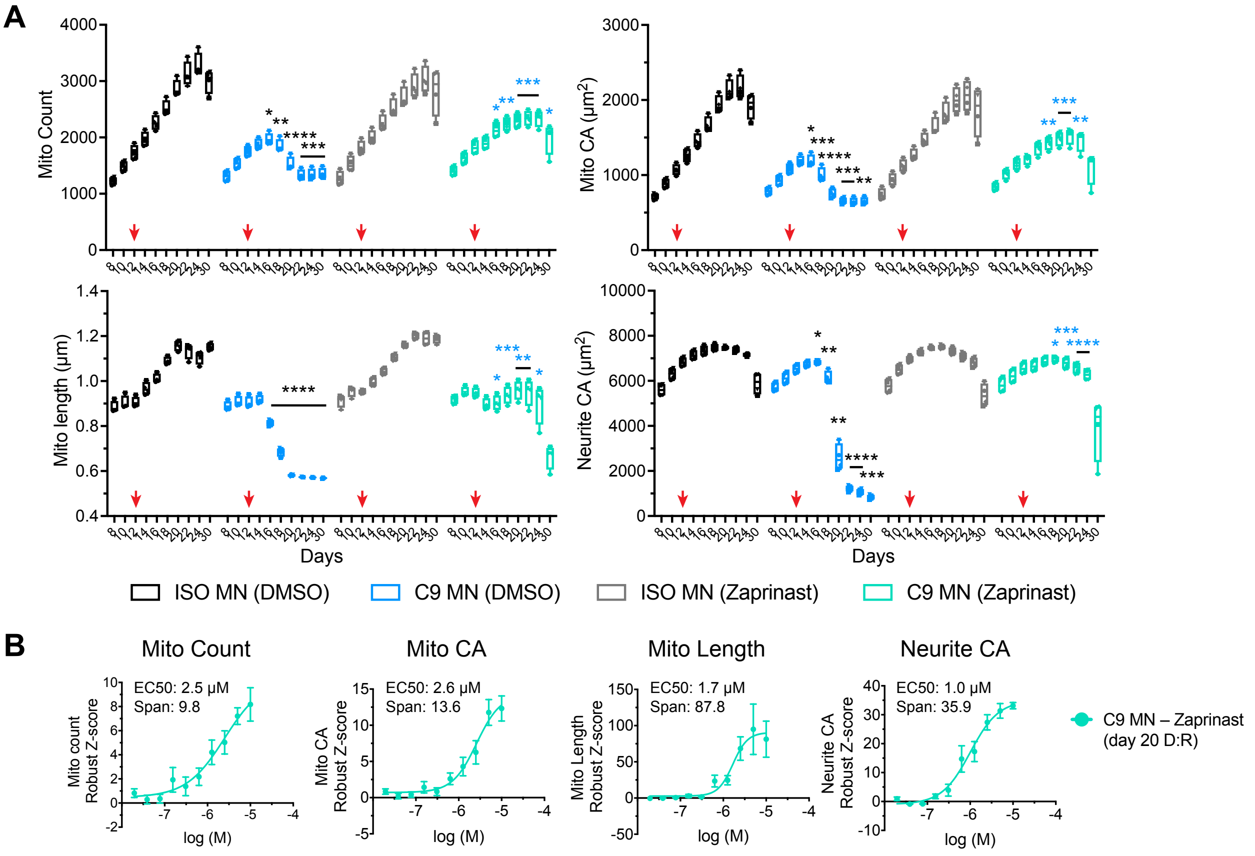

Fig. S9. Time course and D:R experiments for Zaprinast in C9 MN.

**Fig. S9: Time course and D:R experiments for Zaprinast (PDE inhibitor) in C9 MN. (A)** Progressive MT (count, CA, length) and neurite (CA) phenotypes in C9 MN compared to the ISO MN across days in culture in the time-course assay treated with single dose of DMSO or Zaprinast. Data are presented as box plots with individual data points (unique symbols) from n = 4 independent time-course experiments. Box plots show the median (center line), the 25th to 75th percentiles (box), and minimum and maximum values (whiskers). Within each experiment, the DMSO group includes n = 40 technical replicates, while the Zaprinast group includes n = 5 technical replicates. Red arrows indicate the addition of single dose of DMSO (0.125%) / Zaprinast (5 µM) on day 12. Black asterisks illustrate the significant differences of the C9 MN (DMSO) vs ISO MN (DMSO). Blue asterisks illustrate the significant differences of the C9 MN (Zaprinast) vs C9 MN (DMSO). *p<0.05, **p<0.01, ***p<0.001, ****p<0.0001, Two-way repeated measure ANOVA with Tukey's multiple comparisons test. **(B)** Zaprinast D:R data for MT count, MT CA, MT length and neurite CA at day 20 presented as robust Z-scores using C9 MN relative to in-plate DMSO control wells. Zaprinast, tested at concentrations ranging from 20 nM to 10,000 nM, is an inhibitor of PDE 5, 6, 9, and 11, with IC50 values ranging from 150 nM to 29,000 nM, as well as a mitochondrial pyruvate carrier (MPC) inhibitor, with an $IC_{50}$ value of 321 nM.

### Supplemental Tables:

**Table S1: List of iPSC lines used in this study**

| iPSC name | iPSC line | Primary tissue | Mutation | Disease | Sex | Age at sampling (years) | Ethnicity/Race | Comments | IPSCs obtained from: | iPSC Karyotype | Reference |
|---|---|---|---|---|---|---|---|---|---|---|---|
| i3N | i3 Neurons | Fibroblast | | | Male | 30 | Japanese | Non-edited line available from the Corielle Institute of Medical Technology as GM25256 | Corielle Institute of Medical Technology | Normal | Ref. 115 |
| C9 | CS29iALS-C9n1 | Fibroblast | C9orf72 HRE 6-8 kb (1176 repeats) | ALS | Male | 47 | Unknown | Age of onset: 46 years; Site of onset: lower left extremity. Family History: Maternal diagnosed with ALS; maternal grandfather diagnosed with dementia and parkinsonism. | Cedars Sinai Biomanufacturing Center | Normal | https://csbiomfg.com/cellcollection/ |
| ISO | CS29iALS-C9n1.ISO2 RB4 | | | | Male | 47 | Unknown | Isogenic control clones were generated from the C9orf72 iPSCs using the CRISPR-Cas9 complex targeting the regions immediately 5' and 3' of the hexanuclotide repeat (HRE) expansion. | Cedars Sinai Biomanufacturing Center | Normal | https://csbiomfg.com/cellcollection/ |
| BS6 | Becker S6 | Fibroblast | C9orf72 HRE (~750 repeats) | ALS/FTD | Female | 39 | Dutch | Age of onset: 38 years; Behavioral change, language loss, wasting of hand muscles, Dementia, family history of ALS/FTD. | Univ. of Edinburgh (Dr. Chandran's / Dr. Bhuvaneish lab) | Normal | Ref. 114 |
| M211 | M211 R2 | Fibroblast | C9orf72 HRE (~960 repeats) | ALS | Male | 67 | Dutch | Age of onset: 65 years; Lower limb, family history of ALS/FTD. | Univ. of Edinburgh (Dr. Chandran's / Dr. Bhuvaneish lab) | Normal | Ref. 114 |
| 33.1 | 33.1 | Fibroblast | C9orf72 HRE (>620 repeats) | ALS | Male | 65 | Unknown | ALS | Univ. of Dresden (Dr. Jared L. Sterneckert lab) | Normal | Ref. 113 |
| $TDP43^{M337V}$ | KOLF2.1J | Fibroblast | TDP43 M337V | ALS | Male | | White/British | TDP43 M337V mutation was introduced in KOLF2.1J iPSCs by CRISPR-Cas9-based gene editing. | The Jackson Laboratories | Normal | https://www.jax.org/jax-mice-and-services/ipsc |
| REV | KOLF2.1J | | | | Male | | White/British | Isogenic control, TDP43 M337V mutation corrected by CRISPR-Cas9-based gene editing. | The Jackson Laboratories | Normal | https://www.jax.org/jax-mice-and-services/ipsc |
| AG67 | AG25367 | Fibroblast | PSEN1 A246E | AD | Female | 31 | Canadian | Alzheimer's disease. Age of onset: 45 years. | Coriell Repository | Normal | Ref. 52 |
| ISO-AG67 | | | | | Female | 31 | Canadian | Isogenic control, PSEN1 A246E mutation corrected by Synthego via CRISPR-CAS9 gene editing. | Davis Lab | Normal | Ref. 52 |

**Table S2: Dose:Response (D:R) EC50 (µM) and span robust Z-scores for Dipyridamole (DPM) in C9 ALS motor neurons (MN), TDP43$^{M337V}$ ALS MN and PSEN1$^{A246E}$ AD cortical neurons (CN).**

| **C9 ALS MN** | | | | | | | | | | |
|---|---|---|---|---|---|---|---|---|---|---|
| Compound | Dose Response concentration | Days of MN Differentiation | Mito count EC50 (µM) | Mito count span | Mito CA EC50 (µM) | Mito CA span | Mito length EC50 (µM) | Mito length span | Neurite CA EC50 (µM) | Neurite CA span |
| DPM day 12 addition | 20 µM – 156 nM | Day 14 | NR | NR | NR | NR | NR | NR | NR | NR |
| | | Day 16 | **0.93** | 1.7 | **0.95** | 2.5 | **1.34** | 6.3 | **0.57** | 0.9 |
| | | Day 18 | **1.1** | 6.3 | **1.3** | 10.6 | **1.7** | 59.6 | **0.8** | 10.7 |
| | | Day 20 | **1.5** | 9.1 | **1.5** | 12.9 | **1.8** | 59.2 | **1.5** | 24.0 |
| | | Day 22 | **2.1** | 11.7 | **2.4** | 17.4 | **2.6** | 58.8 | **2.1** | 36.0 |
| | | Day 30 | **2.9** | 20.8 | **2.9** | 27.1 | **2.6** | 66.4 | **2.4** | 42.4 |
| DPM day 8 addition | 10 µM – 156 nM | Day 10 | NR | NR | NR | NR | NR | 1.5 | NR | NR |
| | | Day 12 | NR | NR | NR | NR | 0.085 | 2.9 | NR | NR |
| | | Day 14 | **0.140** | 2.0 | **0.114** | 2.6 | **0.128** | 6.9 | **0.084** | 1.6 |
| | | Day 16 | **0.24** | 6.2 | **0.44** | 16.9 | **0.87** | 118.3 | **0.16** | 11.3 |
| | | Day 18 | **0.7** | 7.3 | **1.2** | 12.3 | **1.6** | 102.5 | **1.0** | 56.9 |
| | | Day 20 | **1.6** | 8.8 | **3.5** | 17.2 | **2.7** | 89.4 | **1.9** | 71.7 |
| | | Day 22 | **3.6** | 9.7 | **2.7** | 9.5 | **2.9** | 78.0 | **1.9** | 49.3 |
| | | Day 30 | NR | NR | NR | NR | **5.0** | 41.9 | **4.4** | 117.7 |

| **TDP43$^{M337V}$ ALS MN** | | | | | | | | | | |
|---|---|---|---|---|---|---|---|---|---|---|
| Compound | Dose Response concentration | Days of MN Differentiation | Mito count EC50 (µM) | Mito count span | Mito CA EC50 (µM) | Mito CA span | Mito length EC50 (µM) | Mito length span | Neurite CA EC50 (µM) | Neurite CA span |
| DPM day 10 addition | 10 µM – 156 nM | Day 12 | NR | NR | NR | NR | NR | NR | NR | NR |
| | | Day 14 | 1.8 | 2.1 | 0.8 | 1.4 | NR | NR | NR | NR |
| | | Day 16 | **1.20** | 3.6 | **1.20** | 3.5 | **0.86** | 4.0 | **0.65** | 2.2 |
| | | Day 18 | **1.2** | 6.9 | **1.4** | 7.0 | **1.4** | 9.9 | **0.84** | 9.3 |
| | | Day 20 | **1.5** | 8.6 | **1.5** | 10.3 | **1.9** | 36.8 | **1.3** | 29.5 |
| | | Day 22 | **1.8** | 10.1 | **1.8** | 9.3 | **2.5** | 62.4 | **1.7** | 45.2 |
| | | Day 30 | **2.5** | 21.9 | **2.9** | 10.6 | **3.1** | 24.6 | **2.7** | 224.7 |

| **AG67 CN (PSEN1$^{A246E}$)** | | | | | | | | | | |
|---|---|---|---|---|---|---|---|---|---|---|
| Compound | Dose Response concentration | Days of CN Differentiation | Mito count EC50 (µM) | Mito count span | Mito CA EC50 (µM) | Mito CA span | Mito length EC50 (µM) | Mito length span | Neurite CA EC50 (µM) | Neurite CA span |
| DPM day 10 addition | 10 µM – 20 nM | Day 12 | **8.4** | 4.5 | **4.6** | 2.2 | **12.0** | 6.0 | NR | NR |
| | | Day 14 | **4.5** | 2.8 | **4.5** | 2.9 | NR | NR | NR | NR |
| | | Day 16 | **4.8** | 3.5 | **4.2** | 2.8 | NR | NR | NR | NR |
| | | Day 18 | **5.0** | 5.2 | **5.1** | 4.5 | NR | NR | NR | NR |
| | | Day 20 | NR | NR | **6.8** | 10.1 | **3.6** | 12.9 | NR | NR |
| | | Day 22 | NR | 17.2 | **2.9** | 6.7 | **2.9** | 23.9 | **0.79** | 1.5 |
| | | Day 24 | **7.3** | 9.7 | **2.3** | 8.1 | **2.3** | 41.0 | **0.69** | 3.1 |
| | | Day 26 | **4.4** | 12.3 | **3.2** | 13.4 | NR | NR | **0.59** | 14.3 |
| | | Day 28 | **8.6** | 21.2 | **7.3** | 25.4 | NR | NR | **0.89** | 26.4 |
| | | Day 30 | **5.3** | 17.4 | NR | NR | NR | NR | **1.3** | 21.1 |

NR: not reliable

Potency is determined by EC50, while efficacy is measured by span. Robust Z-scores ≥2 or ≤-2 are considered significant.

| Table S3: Span robust Z-scores for the MT/neurite parameters at different days for the PDE inhibitors tested. | | | | | | | |
|---|---|---|---|---|---|---|---|
| Days of MN Differentiation | Phosphodiesterase (PDE) Inhibitors | Type of PDE Inhibitor | Dose Response concentration | Mito count span Robust Z-score | Mito CA span Robust Z-score | Mito length span Robust Z-score | Neurite CA span Robust Z-score |
| Day 14 | Theophylline | non-selective | 100µM – 2.5 µM | NR | NR | 0.5 | NR |
| | Pentoxifylline | non-selective | 100µM – 2.5 µM | NR | NR | NR | 0.6 |
| | Ibudilast | non-selective | 5 µM – 39 nM | 0.7 | 0.5 | NR | NR |
| | Zaprinast | non-selective | 10 µM – 20 nM | NR | 0.8 | **4.6** | 0.4 |
| | Sildenafil | PDE5 | 40 µM – 625 nM | 1.0 | 0.9 | NR | 1.2 |
| | Vardenafil | PDE5, PDE6 | 1.25 µM – 20 nM | 1.0 | 1.1 | NR | 0.8 |
| | Tadalafil | PDE5, PDE11 | 1.25 µM – 20 nM | NR | NR | 0.5 | 0.0 |
| | PF-04957325 | PDE8 | 1.25 µM – 20 nM | 0.8 | 0.7 | 0.7 | 0.6 |
| | Irsenontrine (E2027) | PDE9 | 10 µM – 20 nM | NR | 0.2 | 0.9 | 0.2 |
| | Osoresnontrine (BI-409306) | PDE9 | 1.25 µM – 20 nM | 0.8 | 0.3 | 0.8 | NR |
| | Mardepodect (PF-2545920 | PDE10 | 1.25 µM – 20 nM | NR | NR | 0.4 | 0.8 |
| | BC11-38 | PDE11 | 1.25 µM – 20 nM | NR | NR | 0.3 | NR |
| Day 16 | Theophylline | non-selective | 100µM – 2.5 µM | 0.4 | 0.7 | 1.6 | NR |
| | Pentoxifylline | non-selective | 100µM – 2.5 µM | NR | NR | NR | NR |
| | Ibudilast | non-selective | 5 µM – 39 nM | NR | NR | 1.3 | 0.3 |
| | Zaprinast | non-selective | 10 µM – 20 nM | **1.9** | **2.3** | **2.4** | **0.7** |
| | Sildenafil | PDE5 | 40 µM – 625 nM | 0.8 | 0.9 | NR | 0.9 |
| | Vardenafil | PDE5, PDE6 | 1.25 µM – 20 nM | 0.7 | NR | NR | NR |
| | Tadalafil | PDE5, PDE11 | 1.25 µM – 20 nM | 0.1 | NR | 0.2 | NR |
| | PF-04957325 | PDE8 | 1.25 µM – 20 nM | 0.4 | 0.4 | NR | NR |
| | Irsenontrine (E2027) | PDE9 | 10 µM – 20 nM | 0.4 | 0.6 | 0.4 | NR |
| | Osoresnontrine (BI-409306) | PDE9 | 1.25 µM – 20 nM | 0.4 | 0.6 | 0.7 | 0.6 |
| | Mardepodect (PF-2545920 | PDE10 | 1.25 µM – 20 nM | NR | NR | NR | NR |
| | BC11-38 | PDE11 | 1.25 µM – 20 nM | 0.7 | 0.6 | 0.8 | 0.5 |
| Day 18 | Theophylline | non-selective | 100µM – 2.5 µM | 0.9 | NR | 2.0 | 1.6 |
| | Pentoxifylline | non-selective | 100µM – 2.5 µM | 0.9 | 0.6 | 1.0 | NR |
| | Ibudilast | non-selective | 5 µM – 39 nM | NR | NR | 0.3 | 0.3 |
| | Zaprinast | non-selective | 10 µM – 20 nM | **8.9** | **11.5** | **30.2** | **8.8** |
| | Sildenafil | PDE5 | 40 µM – 625 nM | 1.5 | 1.4 | NR | NR |
| | Vardenafil | PDE5, PDE6 | 1.25 µM – 20 nM | NR | NR | NR | NR |
| | Tadalafil | PDE5, PDE11 | 1.25 µM – 20 nM | 0.3 | 0.3 | 0.9 | 0.3 |
| | PF-04957325 | PDE8 | 1.25 µM – 20 nM | NR | NR | NR | NR |
| | Irsenontrine (E2027) | PDE9 | 10 µM – 20 nM | 0.6 | 0.7 | 2.2 | NR |
| | Osoresnontrine (BI-409306) | PDE9 | 1.25 µM – 20 nM | 0.6 | 0.4 | NR | 1.0 |
| | Mardepodect (PF-2545920 | PDE10 | 1.25 µM – 20 nM | NR | 1.6 | NR | 1.0 |
| | BC11-38 | PDE11 | 1.25 µM – 20 nM | NR | NR | NR | NR |

| | | | | | | | |
|---|---|---|---|---|---|---|---|
| Day 20 | Theophylline | non-selective | 100µM – 2.5 µM | NR | NR | 1.0 | NR |
| | Pentoxifylline | non-selective | 100µM – 2.5 µM | NR | NR | NR | NR |
| | Ibudilast | non-selective | 5 µM – 39 nM | NR | NR | 0.2 | NR |
| | Zaprinast | non-selective | 10 µM – 20 nM | **9.8** | **13.6** | **87.8** | **36.0** |
| | Sildenafil | PDE5 | 40 µM – 625 nM | NR | NR | NR | NR |
| | Vardenafil | PDE5, PDE6 | 1.25 µM – 20 nM | NR | NR | NR | 0.3 |
| | Tadalafil | PDE5, PDE11 | 1.25 µM – 20 nM | 0.5 | 0.5 | NR | NR |
| | PF-04957325 | PDE8 | 1.25 µM – 20 nM | NR | NR | NR | NR |
| | Irsenontrine (E2027) | PDE9 | 10 µM – 20 nM | NR | NR | 3.2 | NR |
| | Osoresnontrine (BI-409306) | PDE9 | 1.25 µM – 20 nM | 0.4 | 0.4 | NR | 0.5 |
| | Mardepodect (PF-2545920 | PDE10 | 1.25 µM – 20 nM | NR | NR | NR | NR |
| | BC11-38 | PDE11 | 1.25 µM – 20 nM | NR | NR | NR | NR |
| Day 22 | Theophylline | non-selective | 100µM – 2.5 µM | NR | NR | NR | NR |
| | Pentoxifylline | non-selective | 100µM – 2.5 µM | NR | NR | NR | NR |
| | Ibudilast | non-selective | 5 µM – 39 nM | NR | NR | NR | NR |
| | Zaprinast | non-selective | 10 µM – 20 nM | **7.8** | **9.6** | **44.4** | **34.5** |
| | Sildenafil | PDE5 | 40 µM – 625 nM | NR | NR | NR | NR |
| | Vardenafil | PDE5, PDE6 | 1.25 µM – 20 nM | NR | NR | NR | NR |
| | Tadalafil | PDE5, PDE11 | 1.25 µM – 20 nM | NR | NR | NR | NR |
| | PF-04957325 | PDE8 | 1.25 µM – 20 nM | NR | NR | NR | NR |
| | Irsenontrine (E2027) | PDE9 | 10 µM – 20 nM | NR | NR | NR | NR |
| | Osoresnontrine (BI-409306) | PDE9 | 1.25 µM – 20 nM | NR | NR | NR | NR |
| | Mardepodect (PF-2545920 | PDE10 | 1.25 µM – 20 nM | NR | NR | NR | NR |
| | BC11-38 | PDE11 | 1.25 µM – 20 nM | NR | NR | NR | NR |
| Day 24 | Theophylline | non-selective | 100µM – 2.5 µM | NR | NR | NR | NR |
| | Pentoxifylline | non-selective | 100µM – 2.5 µM | NR | NR | NR | NR |
| | Ibudilast | non-selective | 5 µM – 39 nM | NR | NR | NR | NR |
| | Zaprinast | non-selective | 10 µM – 20 nM | **4.1** | **4.4** | **14.2** | **25.9** |
| | Sildenafil | PDE5 | 40 µM – 625 nM | NR | NR | NR | NR |
| | Vardenafil | PDE5, PDE6 | 1.25 µM – 20 nM | NR | NR | NR | NR |
| | Tadalafil | PDE5, PDE11 | 1.25 µM – 20 nM | NR | NR | NR | NR |
| | PF-04957325 | PDE8 | 1.25 µM – 20 nM | NR | NR | NR | NR |
| | Irsenontrine (E2027) | PDE9 | 10 µM – 20 nM | NR | NR | NR | NR |
| | Osoresnontrine (BI-409306) | PDE9 | 1.25 µM – 20 nM | NR | NR | NR | NR |
| | Mardepodect (PF-2545920 | PDE10 | 1.25 µM – 20 nM | NR | NR | NR | NR |
| | BC11-38 | PDE11 | 1.25 µM – 20 nM | NR | NR | NR | NR |
| | NR: not reliable | | | | | | |

Span (efficacy) is measure by robust Z-score relative to DMSO-only C9 MN-containing wells. Robust Z-scores ≥2 or ≤-2 are considered significant.

**Table S4: Span robust Z-scores for the MT/neurite parameters at different days for the ENT1/ENT2 inhibitors tested.**

| Days of MN Differentiation | ENT1/ENT2 Inhibitors | Dose Response concentration | Mito count span Robust Z-score | Mito CA span Robust Z-score | Mito length span Robust Z-score | Neurite CA span Robust Z-score |
|---|---|---|---|---|---|---|
| Day 14 | NBTI | 40 nM - 0.313 nM | 0.6 | 0.5 | NR | 0.1 |
| | Draflazine | 1.25 µM – 20 nM | 1.1 | 1.1 | 0.3 | 0.9 |
| | JMF 1907 | 1.25 µM – 20 nM | 1.3 | NR | NR | 0.4 |
| Day 16 | NBTI | 40 nM - 0.313 nM | 0.2 | 0.3 | 0.8 | 0.0 |
| | Draflazine | 1.25 µM – 20 nM | 0.3 | NR | 0.9 | NR |
| | JMF 1907 | 1.25 µM – 20 nM | NR | NR | NR | 1.2 |
| Day 18 | NBTI | 40 nM - 0.313 nM | 0.7 | NR | 0.7 | 0.8 |
| | Draflazine | 1.25 µM – 20 nM | NR | NR | 0.7 | NR |
| | JMF 1907 | 1.25 µM – 20 nM | NR | NR | NR | NR |

NR: not reliable

Span (efficacy) is measure by robust Z-score relative to DMSO-only C9 MN-containing wells. Robust Z-scores ≥2 or ≤-2 are considered significant.

**Table S5: Span robust Z-scores for the MT/neurite parameters at different days for the SREBP inhibitors tested.**

| Days of MN Differentiation | SREBP Inhibitors | Dose Response concentration | Mito count span Robust Z-score | Mito CA span Robust Z-score | Mito length span Robust Z-score | Neurite CA span Robust Z-score |
|---|---|---|---|---|---|---|
| Day 14 | Fatostatin | 10 µM – 156 nM | NR | NR | 1.2 | NR |
| | Betulin | 10 µM – 156 nM | NR | NR | NR | NR |
| | PF429242 | 10 µM – 156 nM | NR | 0.5 | NR | 0.6 |
| Day 16 | Fatostatin | 10 µM – 156 nM | NR | NR | 1.3 | 0.6 |
| | Betulin | 10 µM – 156 nM | 1.2 | 1.3 | NR | NR |
| | PF429242 | 10 µM – 156 nM | 0.9 | 0.8 | 1.7 | NR |
| Day 18 | Fatostatin | 10 µM – 156 nM | NR | NR | NR | NR |
| | Betulin | 10 µM – 156 nM | NR | NR | NR | NR |
| | PF429242 | 10 µM – 156 nM | NR | NR | NR | 0.3 |

NR: not reliable

Span (efficacy) is measure by robust Z-score relative to DMSO-only C9 MN-containing wells. Robust Z-scores ≥2 or ≤-2 are considered significant.

| Table S6. Resources and reagents used. | | |
|---|---|---|
| **Reagent or Resource** | **Source** | **Identifier** |
| **Antibodies** | | |
| Mouse anti-HB9 (MNR2/HB9/Mnx1) | DSHB | 81.5C10 |
| Mouse anti-NFH (SMI32) | Fisher Scientific (BioLegend) | 801701 |
| Rabbit anti-β-Tubulin III (Tuj1) | Millipore-Sigma | T2200 |
| Goat anti-ChAT | Millipore-Sigma | AB144P |
| Mouse anti-Tau (clone tau46) | Millipore-Sigma | T9450 |
| Rabbit anti-MAP2 | Abcam | ab32454 |
| Rabbit anti-Oct4 | Abcam | ab18976 |
| Mouse anti-SOX2 | Abcam | ab79351 |
| MPC1 (D2L9I) Rabbit Monoclonal Antibody | Cell Signaling Technology | 14462 |
| MPC2 (D4I7G) Rabbit Monoclonal Antibody | Cell Signaling Technology | 46141 |
| Vinculin (E1E9V) Rabbit Monoclonal Antibody | Cell Signaling Technology | 13901 |
| Alexa Fluor 568 Donkey anti-Rabbit IgG (H+L) | ThermoFisher Scientific | A10042 |
| Alexa Fluor 488 donkey anti-Rabbit IgG (H+L) | ThermoFisher Scientific | A21206 |
| Alexa Fluor 488 Donkey anti-Mouse IgG | ThermoFisher Scientific | A21202 |
| Alexa Fluor 488 Donkey anti-Goat IgG (H+L) | ThermoFisher Scientific | A11055 |
| Alexa Fluor 568 Donkey anti Goat IgG (H+L) | ThermoFisher Scientific | A11057 |
| IRDye 800CW goat anti-rabbit IgG | LI-COR | 926–32211 |
| **Bacterial and virus strains** | | |
| pCAG-mtTagGFP2-2A-mScarlet plasmid was packaged into a lentivirus | SignaGen Laboratories | Ref. 52 |
| AAV8-TBG-Cre adeno-associated virus serotype 8 | Vector Biolabs | VB1724 |
| AAV8-TBG-eGFP | Vector Biolabs | VB1743 |
| **Chemicals, peptides, and recombinant proteins** | | |
| mTeSR1 kit (Basal medium + 5x Supplement) | StemCell Technologies | 85850 |
| mTeSR Plus kit (Basal medium + 5x Supplement) | StemCell Technologies | 100-0276 |
| Dulbecco's Phosphate Buffered Saline (DPBS without $Ca2^{+}/Mg2^{+}$) | ThermoFisher Scientific | 14190144 |
| Matrigel (Growth Factor Reduced) | Corning | 354230 |
| StemFlex Medium | ThermoFisher Scientific | A3349401 |
| Synthemax II-SC Substrate | Fisher Scientific | 7201611 |
| DMEM/F-12 with GlutaMAX™ supplement | ThermoFisher Scientific | 10565018 |
| StemPro Accutase Cell Dissociation Reagent | ThermoFisher Scientific | A1110501 |
| ReLeSR | StemCell Technologies | 100-0483 |
| UltraPure 0.5M EDTA, pH 8.0 | ThermoFisher Scientific | 15575020 |
| CryoStor CS10 | StemCell Technologies | 100-1061 |
| Alt-R S.p. HiFi Cas9 Nuclease V3 | Integrated DNA Technologies | 1081060 |
| Lipofectamine Stem transfection reagent | ThermoFisher Scientific | STEM00003 |
| Opti-MEM™ I Reduced Serum Medium | ThermoFisher Scientific | 31985062 |
| Y-27632 (Dihydrochloride) | StemCell Technologies | 72302 |
| Blasticidin | InvivoGen | ant-bl-1 |
| QuickExtract DNA Extract Solution | Biosearch Technologies | QE09050 |
| Platinum SuperFi II PCR Master Mix | ThermoFisher Scientific | 12368010 |
| DMEM/F12 with HEPES and glutamine | ThermoFisher Scientific | 11330032 |
| B-27™ Plus Neuronal Culture System (has Neurobasal plus media and B27 plus supplement) | ThermoFisher Scientific | A3653401 |
| GlutaMAX supplement | ThermoFisher Scientific | 35050061 |
| MEM Non-Essential Amino Acids Solution (100X) | ThermoFisher Scientific | 11140050 |
| CultureOne supplement | ThermoFisher Scientific | A3320201 |
| Doxycycline hyclate | Millipore-Sigma | D9891 |
| γ-Secretase Inhibitor XXI, Compound E | Millipore-Sigma | 565790 |
| BamBanker Cell Freezing Media, BB02 | Fisher Scientific | 50999555 |
| Laminin mouse protein | ThermoFisher Scientific | 23017015 |
| Poly-D-Lysine (0.1 mg/mL, 50,000–150,000 Daltons) | ThermoFisher Scientific | A3890401 |
| Penicillin-Streptomycin (5,000 U/mL) | ThermoFisher Scientific | 15070063 |
| Recombinant Human/Murine/Rat BDNF | PeproTech | 450-02-10µg |
| Recombinant human NT-3 | PeproTech | 450-03-10µg |
| Recombinant human GDNF | PeproTech | 450-10-10µg |
| 16% paraformaldehyde | Electron Microscopy Sciences | 15710 |
| Triton X-100 | Millipore-Sigma | T8787 |
| Bovine serum albumin | Millipore-Sigma | A8806 |
| Calcein, AM, cell-permeant dye | ThermoFisher Scientific | C3100MP |
| Oligomycin from Streptomyces diastatochromogenes | Millipore-Sigma | O4876 |
| Carbonyl cyanide 4-(trifluoromethoxy)phenylhydrazone (FCCP) | Millipore-Sigma | C2920 |
| Rotenone | Millipore-Sigma | R8875 |
| Antimycin A from Streptomyces sp. | Millipore-Sigma | A8674 |
| Seahorse XFe96/XF Pro FluxPak | Agilent | Part# 103792-100 |
| Seahorse XF DMEM medium, pH 7.4 | Agilent | Part# 103575-100 |
| Seahorse XF 100 mM pyruvate solution | Agilent | Part# 103578-100 |
| Seahorse XF 1.0 M glucose solution | Agilent | Part# 103577-100 |
| Seahorse XF 200 mM glutamine solution | Agilent | Part# 103579-100 |
| 384 well PDL coated black plates | Aurora Microplates | ABE2-0201B-PDL |
| 96 well µClear black plates | Greiner | 655090 |

| | | |
|---|---|---|
| Dimethyl sulfoxide (DMSO) | Millipore-Sigma | D2650 |
| Dipyridamole | Millipore-Sigma | D9766 |
| NBTI | Millipore-Sigma | N2255 |
| Draflazine | Millipore-Sigma | JN0006 |
| JMF 1907 | Cayman chemicals | 28414 |
| Theophylline anhydrous | Millipore-Sigma | T1633 |
| Pentoxifylline | Millipore-Sigma | P1784 |
| Ibudilast | Millipore-Sigma | I0157 |
| Sildenafil Citrate | Millipore-Sigma | SML3033 |
| Vardenafil hydrochloride trihydrate | MedChem Express | HY-B0442B |
| Tadalafil | Millipore-Sigma | SML1877 |
| PF-04957325 | MedChem Express | HY-15426 |
| Irsenontrine (E2027) | MedChem Express | HY-132821 |
| Osoresnontrine (BI-409306) | MedChem Express | HY-112831 |
| Mardepodect hydrochloride (PF-2545920 hydrochloride) | Millipore-Sigma | PZ0351 |
| BC11-38 | MedChem Express | HY-108618 |
| Fatostatin | MedChem Express | HY-14452 |
| Betulin | MedChem Express | HY- N0083 |
| PF429242 dihydrochloride | MedChem Express | HY- 13447A |
| Mannitol | Millipore Sigma | M4125-500G |
| Sucrose | Millipore Sigma | S0389-500G |
| HEPES | ThermoFisher Scientific | 15630080 |
| EGTA | Fisher Scientific | 50255956 |
| KOH | Millipore Sigma | 484016-1KG |
| BSA | ThermoFisher Scientific | B14 |
| KCl | Millipore Sigma | P9541-500G |
| Pierce BCA protein assay | Fisher Scientific | PI23225 |
| UK-5099 | Fisher Scientific | 502260165 |
| Pyruvic acid sodium salt (2-C14) | Revvity Health Sciences | NEC256050UC |
| Ultima gold scintillation fluid | Revvity Health Sciences | 6013321 |
| RIPA buffer | Cell Signaling Technology | 9806S |
| RIPA buffer | ThermoFisher Scientific | 89900 |
| Malate | Sigma | M1000 |
| ADP | Merck | 117105 |
| Cytochrome C | Sigma | C7752 |
| Sodium Pyruvate | Sigma | P2256 |
| **Critical commercial assays/kits** | | |
| QIAprep Spin Miniprep Kit | Qiagen | 27106 |
| PureLink HiPure Plasmid Filter Maxiprep Kit | ThermoFisher Scientific | K210017 |
| RealTime-Glo MT Cell Viability Assay | Promega | G9711 |
| **Experimental models: organisms/strains** | | |
| Mouse: C57BL/6J | The Jackson Laboratory | 664 |
| Mpc2 Flox mice | McCommis et al, 2015 Ref. 111 | N/A |
| **Oligonucleotides** | | |
| custom sgRNA ATGTTGGAAGGATGAGGAAA (targeting the human CLYBL intragenic safe harbor locus (between exons 2 and 3) | Synthego | N/A |
| CLYBL WT F: TGACTAAACACTGTGCCCCA | Integrated DNA Technologies | N/A |
| CLYBL WT R: AGGCAGGATGAATTGGTGGA | Integrated DNA Technologies | N/A |
| hNIL CLYBL 5' Junction F: CAGACAAGTCAGTAGGGCCA | Integrated DNA Technologies | N/A |
| hNIL CLYBL 5' Junction R: AGAAGACTTCCTCTGCCCTC | Integrated DNA Technologies | N/A |
| hNIL CLYBL 3' Junction F: CACCAGCAACCTGACGTTTT | Integrated DNA Technologies | N/A |
| hNIL CLYBL 3' Junction R: TTTTATAGGCGCCCACCGTA | Integrated DNA Technologies | N/A |
| **Recombinant DNA** | | |
| pCAG-mtTagGFP2-2A-mScarlet plasmid | Davis lab | Ref. 52 |
| CLYBL-TO-hNIL-BSD-mApple | a gift from Michael Ward | Addgene plasmid # 124230 ; http://n2t.net/addgene:124230 ; RRID:Addgene_124230 |
| pCE-mp53DD | a gift from Shinya Yamanaka | Addgene plasmid # 41856 ; http://n2t.net/addgene:41856 ; RRID:Addgene_41856 |
| **Software and algorithms** | | |
| MetaXpress software v6.5.4.532 | Molecular Devices | https://www.moleculardevices.com |
| GraphPad Prism v10.4.1 | GraphPad Software | https://www.graphpad.com/ |
| Fiji (ImageJ2) v2.16.0 | | https://imagej.net/software/fiji/downloads |
| InCell Developer 1.9.2 software | GE Healthcare Technologies | |
| SnapGene v7.1.2 | SnapGene | https://www.snapgene.com/ |
| JMP v.15 | JMP Statistical Discovery | https://www.jmp.com/ |
| DatLab | Oroboros Instruments | https://www.mitofit.org/index.php/DatLab_7 |
| **Other** | | |
| EVOS XL Core Imaging System | ThermoFisher Scientific | https://www.thermofisher.com |
| Image Xpress Micro Confocal High-Content Imaging System | Molecular Devices | https://www.moleculardevices.com |
| InCell 6000 analyzer | GE Healthcare Technologies | |
| CLARIOstar microplate reader | BMG Labtech | https://www.bmglabtech.com |
| Seahorse XFe96 analyzer | Agilent | https://www.agilent.com |
| Oroboros Oxygraph-2K (O2K) system | Oroboros Instruments | https://www.oroboros.at 10203-03 |
| Overhead stirrer | Glas-Col | www.glascol.com |
| Hidex scintillation counter | Hidex (Turku, Finland) | www.hidex.com |
| Tissuelyser | Qiagen | https://www.qiagen.com |
| Fluorescence imaging system (LI-COR) | LI-COR | https://www.licorbio.com |

**Supplemental Movies**

**Movie S1:** C9 MNs at day 110 treated with DPM at day 12, illustrating the trafficking of MT in the C9 neurites.

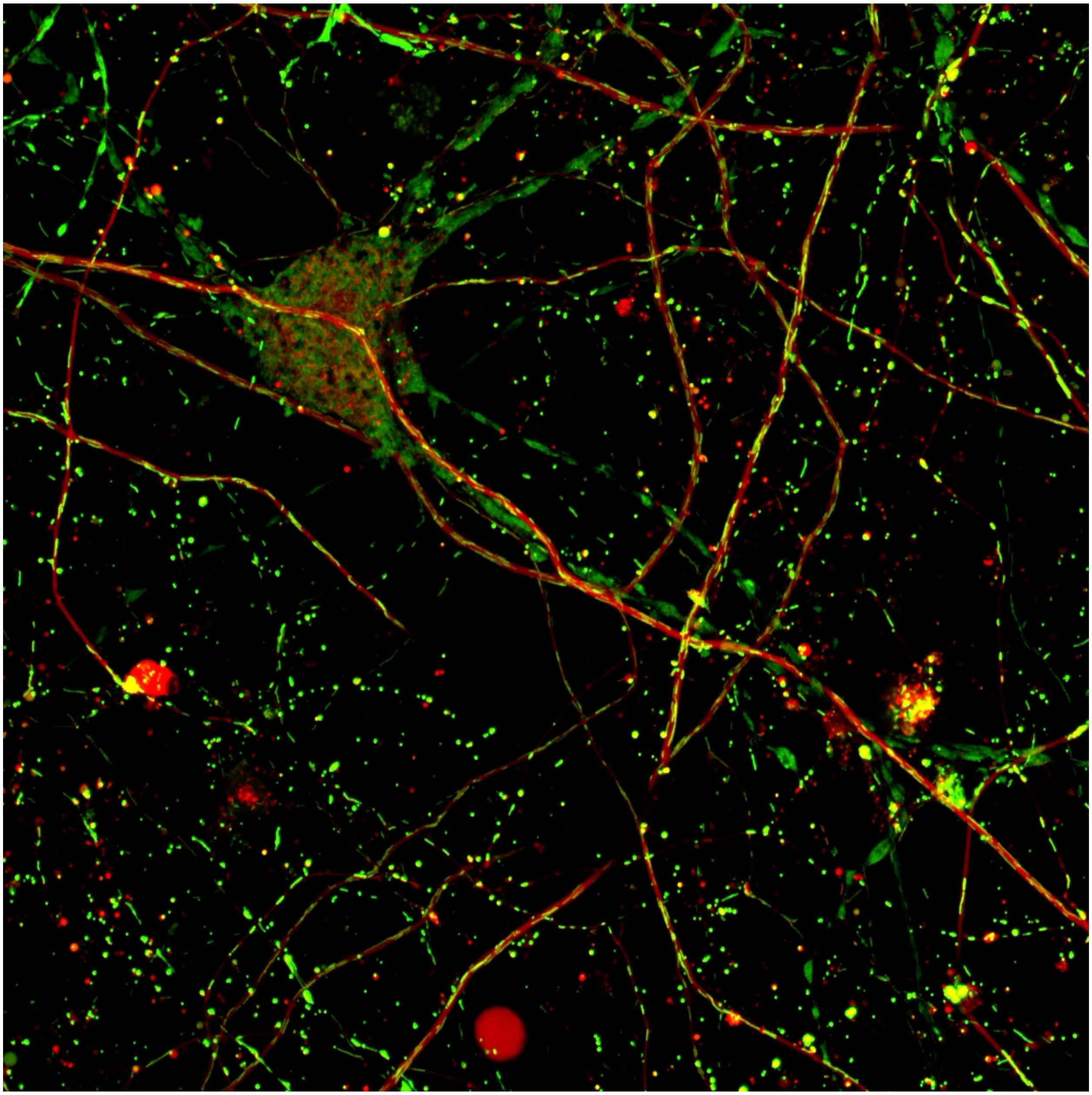
